# Structural basis for assembly of TRAPPII complex and specific activation of GTPase Ypt31/32

**DOI:** 10.1101/2021.11.11.468031

**Authors:** Chenchen Mi, Li Zhang, Shan Sun, Guoqiang Huang, Guangcan Shao, Fan Yang, Xin You, Meng-Qiu Dong, Sen-Fang Sui

**Affiliations:** State Key Laboratory of Membrane Biology, Beijing Advanced Innovation Center for Structural Biology, Beijing Frontier Research Center for Biological Structure, School of Life Sciences, Tsinghua University, 100084, Beijing, China; National Institute of Biological Sciences, Beijing 102206, China; Department of Biology, Southern University of Science and Technology, Shenzhen, Guangdong 518055, China

## Abstract

Transport protein particle (TRAPP) complexes belong to the multiprotein tethering complex and have three forms- TRAPPI, TRAPPII and TRAPPIII, which share a core of six TRAPPI proteins. TRAPPII facilitates intra-Golgi and endosome-to-Golgi transports by activating GTPase Ypt31/Ypt32 as the guanine nucleotide exchange factor (GEF) in yeast. Here we present cryo-EM structures of yeast TRAPPII in apo and Ypt32-bound states. All the structures show a dimeric architecture assembled by two triangle shaped monomers, while the monomer in the apo structure exhibits both open and closed conformations, and the monomer in the Ypt32-bound form only captures the closed conformation. Located in the interior of the monomer, Ypt32 binds with both TRAPPI and Trs120 via its nucleotide binding domain and binds with Trs31 of TRAPPI via its hypervariable domain. Combined with functional analysis, the structures provide insights into the assembly of TRAPPII and the mechanism of the specific activation of Ypt31/Ypt32 by TRAPPII.

**One Sentence Summary:** Structures of TRAPPII in different states reveal the mechanism of the specific activation of Ypt32 by TRAPPII.

## Main Text

In eukaryotic cells, vesicles act as the cargo carriers by transporting proteins, lipids and other materials between various membrane-bound compartments (*1*). Vesicle transport is a successive process. Each step along the pathway, from the vesicle budding at the donor compartment, via vesicle transporting in cell, to the tethering and fusion with the membrane of the acceptor compartment, is precisely controlled by corresponding factors (*2*). Tethering refers to the initial interaction between a vesicle and its target membrane (*2*) and promotes the organization of the soluble N-ethylmaleimide–sensitive factor attachment protein receptor (SNARE) proteins in vesicle fusion (*3*), and thus is a crucial step in determining the specificity of vesicle trafficking (*2*). The tethering process is highly regulated by the tethering factors, including the conserved GTPases of the Ypt/Rab family, their guanine nucleotide exchange factors (GEFs), and their downstream effectors (*4–7*). How the different types of tethering factors work together to achieve the overall specificity of the tethering process remains to be elucidated.

Transport protein particle (TRAPP) complexes belong to the multisubunit protein complexes and are highly conserved family of proteins found in all eukaryotes, from yeast to humans. TRAPP complexes from yeast are the well-studied members of this family and have three forms-TRAPPI, TRAPPII and TRAPPIII, which function in various vesicle trafficking pathways (*8*). TRAPPI mediates transport from ER to cis-Golgi by tethering COPII-coated vesicles (*5, 9*); TRAPPII interacts with COPI-coated vesicles and aids in intra-Golgi and endosome-to-Golgi transport (*6, 10, 11*); TRAPPIII plays a unique role in autophagy (*12*). The TRAPP complexes attract particular attention in the vesicle transport because they have been found acting as GEFs to play an essential role in catalyzing nucleotide exchange for Ypt/Rab GTPasae. All three complexes share a core of six proteins (Bet3, Bet5, Trs20, Trs23, Trs31, Trs33) that make up TRAPPI (*13*). TRAPPII contains four additional proteins (Trs120, Trs130, Trs65, Tca17) (*14*) and TRAPPIII has one additional Trs85 (*15*). A common core suggests common functions with all complexes acting as GEFs. TRAPPI and TRAPPIII are specific to Ypt1 (*16–18*), whereas TRAPPII activates Ypt31/32 (*11, 19, 20*). So far, the detailed mechanisms underlying the transition from TRAPPI and TRAPPIII’s GEF activity for Ypt1 to TRAPPII’s GEF activity for Ypt31/32 and the functions of the specific subunits of TRAPPII in this transition are poorly understood.

Since TRAPPI is the common core of TRAPP complexes, it has been a subject of various in-depth studies on its structure. A combination of X-ray crystallography and single particle electron microscopy (EM) showed an elongated rod-shaped structure of TRAPPI, containing seven subunits (two copies of Bet3 and one copy of each of the other proteins) arranged side by side (*21*). The crystal structure of the yeast TRAPPI core in complex with Ypt1 revealed that four subunits (Bet5p, Trs23 and two Bet3 subunits) interact directly with Ypt1 to stabilize Ypt1 in an open conformation, facilitating nucleotide exchange (*22*). TRAPPII is the largest member in the TRAPP family, and its architecture was proposed based on the negatively stained single-particle EM. The data showed that TRAPPII dimerizes into a three-layered, diamond-shaped structure with two TRAPPI complexes forming the outer layers and the TRAPPII-specific subunits forming the middle layer (*23*). TRAPPI is preserved in TRAPPII, but TRAPPI and TRAPPII activate different Rab substrates: TRAPPI activates Ypt1 and TRAPPII activates Ypt31/32 (*24*). Thus, whether TRAPPII use the same catalytic site as TRAPPI, and how TRAPPII specifically active Ypt31/32, are the problems that perplexes people and there are contradictory results (*11, 17, 25, 26*).

To address these questions and the fundamental mechanism of TRAPPII in vesicle trafficking, we resolved the structures of the intact TRAPPII and its complex with Ypt32 from *Saccharomyces cerevisiae* at average resolutions of 3.71 Å and 3.86 Å, respectively, by single-particle cryo-electron microscopy (cryo-EM). Combination with biochemistry analyses, the structures revealed the detailed interactions between subunits within TRAPPII and between TRAPPII and Ypt32, as well as the dynamic conformations of TRAPPII. Based on these results, the potential working mechanism for TRAPPII in the vesicle trafficking is discussed.

### Overall structure of the TRAPP II complex

The intact TRAPP II complex with the molecular mass of about 1052 kilodalton was purified from a yeast strain containing FLAG-tagged Trs120. SDS-PAGE analysis and mass spectrometry (MS) indicated that all 10 different proteins, including six TRAPPI proteins (Bet5p, Bet3p, Trs20p, Trs23p, Trs31p, and Trs33p) and four TRAPPII-specific proteins (Tca17, Trs65p, Trs120p, and Trs130p), are present in the purified TRAPPII complex (figs. S1, A and B). Especially, Tca17, which was not detected in the previously published structure (*23*), appears to be a stoichiometric component in our sample (fig. S1A). Initial reconstruction led to a structure showing an apparent two-fold symmetry (figs.S1, C to E), which is consistent with the reported low-resolution negative staining EM structure that TRAPPII is a dimeric complex (*23*). However, only one half of this reconstruction displayed clear density and the other half exhibited very poor density suggesting its heterogeneity (fig. S1E). To deal with this, we expanded the data set by rotating one monomer (half of the entire dimer) 180° along the C2 symmetry axis so that both monomers are reoriented onto a single position. We then performed classification by applying a mask around the reoriented monomers (fig. S1F). Through this strategy, two distinct conformations of monomer were identified, and 3D refinement yielded the closed and open structures at resolutions of 3.71 Å and 4.15 Å, respectively (figs. S1, F to H, and S2A). The 2:1 ratio of the particles between the closed and open conformation likely suggests the closed conformation is more stable (fig. S1F). We further assigned each monomer back to its original dimeric particle, resulting in the reconstruction of TRAPPII structure in three different states (fig. S1F). State I contains both monomers in closed conformation and State III contains both monomers in open conformation (figs. S1F and S2B). Different from them, one monomer is in open conformation and the other one is in closed conformation in State II (figs. S1F and S2B). Based on these maps, the atomic models of TRAPPII in open and closed conformation were built (figs. S1, I and J).

The overall structure of TRAPPII looks like an arch bridge with about 160 Å height from the side view and has a parallelogram outline from the face view with dimensions of approximate 290 Å by 260 Å (Fig. 1A). The dimeric complex is assembled by two triangle shaped monomers, which are associated with each other through the longest edge composed of Trs120 and Trs65 (Fig. 1A and fig. S2B). TRAPPI and Trs130, which are connected by Tca17, form the middle and shortest edges of the triangle, respectively (Fig. 1A). For monomers in the complex, the major difference between the open conformation and the closed conformation lies in the position of TRAPPI (Fig. 1D). Once Trs120, Trs65 and Trs130 have been superimposed (fig. S2C), the TRAPPI rotates 16.6° towards the Trs120 pivoting around the TRAPPI-Trs120 junction, together with a slight turn (8.9°) on its own axis, from the open conformation to the closed conformation, which leads to the reduce of the angle between TRAPPI and Trs120 by about 10° (from 47.3° to 36.9°) (Fig. 1D). Such differences led to the larger space of the interior of TRAPPII in the open conformation than in the closed conformation.

**Figure 1.**
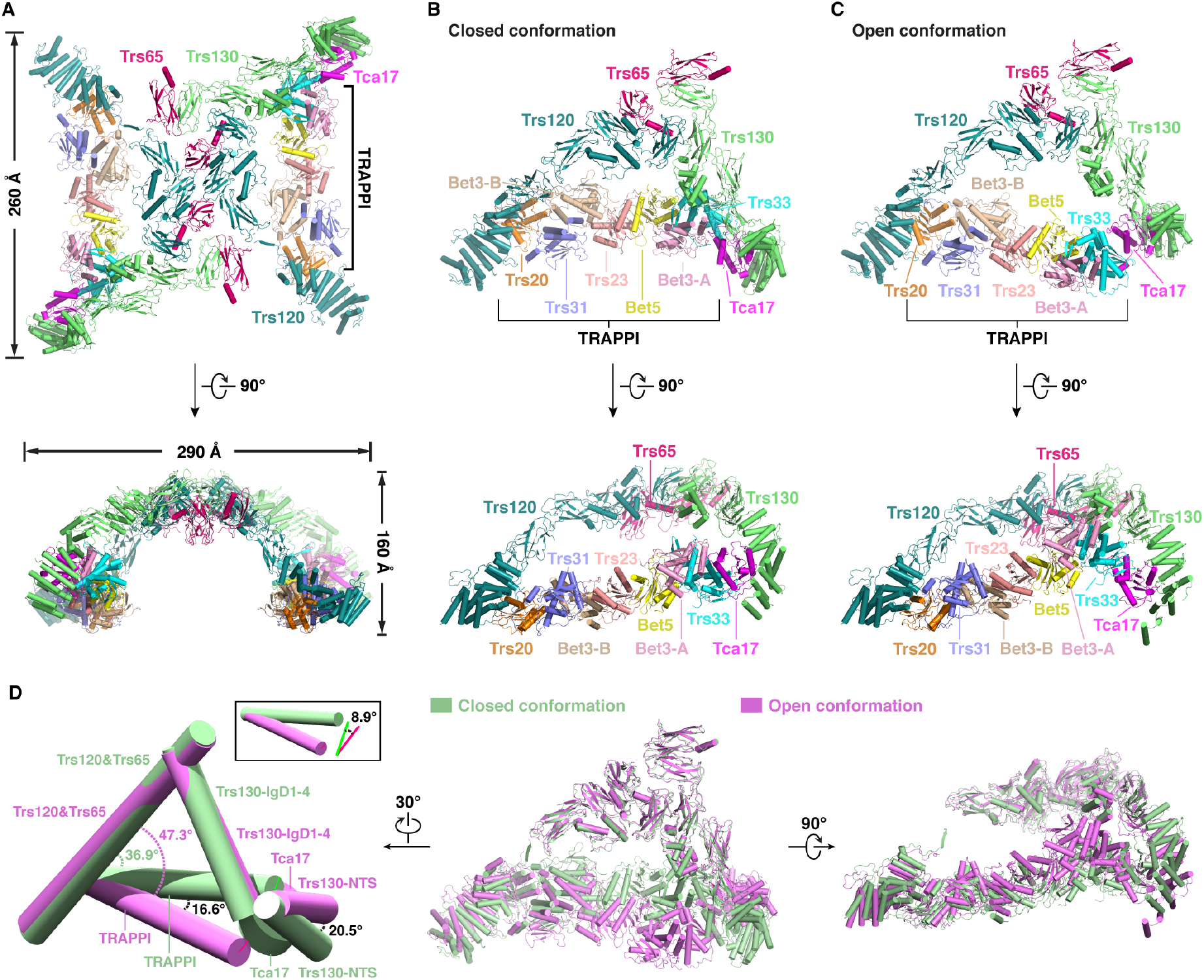
Structures of the yeast TRAPPII complex. **(A)** The overall structure of the intact yeast TRAPPII in State I. **(B)** The structure of the yeast TRAPPII monomer in the closed conformation**. (C)** The structure of the yeast TRAPPII monomer in the open conformation. **(D)** Structural comparison of the closed and the open conformations.

### Structures of TRAPPI and Tca17 in TRAPPII

The structure of TRAPPI was well resolved in our EM map and the complete atomic model was built with all 7 subunits assigned accurately (Figs. 1, B and C, and fig. S2A). TRAPPI looks like a flat rod about 180 Å long and is formed by the arrangement of Trs33, Bet3-A, Bet5, Trs23, Bet3-B, Trs31 and Trs20 in turn (Figs. 1, B and C), which is in agreement with the previously supposed organization of TRAPPI based on the crystal structures and the low-resolution EM structure (*21*). Most of the structures are well superimposed with the existing crystal structures, and some loops that were absent in crystal structures are clearly resolved in our maps (fig. S3A). Besides the interactions between subunits revealed by the crystal structures of subcomplexes, the most striking finding in our cryo-EM structure is that Trs31 extends its long N-terminal loop (residues 25-54) first passing through the hole between loop α1-α2 and the C-terminal end of helix α4 of Bet3-B and then wandering around the surface of Trs20 (fig. S3B), which functions like an arm to hold Trs20 firmly. Compared with the human Trs33, the yeast Trs33 has extra two β strands and one short α helix (residues 87-109) following the first helix (fig. S3C). The two β strands are docked to the groove formed by the helices α3 and α5, and β strand β1 of Bet3-A. The short α helix contacts with the loop α2-α3 of Bet5 (fig. 3C). Together, these interactions could enhance the associations among subunits of TRAPPI, thus making the whole complex more stable and rigid.

After docking TRAPPI into the EM map, we can still observe some clear extra density next to the Trs33 (fig. S2A). We attributed this density to the Tca17 protein. Indeed, the crystal structure of the Tca17 (PDB ID: 3PR6) fits into the density with high confidence (fig. S2D). Thus, Tca17 is located on the opposite end of the TRAPPI rod to Trs20 and interacts with Trs33 in the TRAPPII complex (Figs. 1, B and C). On the other side, Tca17 also interacts with Trs130, which will be described below.

### Structure of Trs120 in TRAPPII

Since Trs120 and Trs130 have the similar overall domain arrangement that is one N-terminal α-solenoid (NTS) domain followed by four Ig-like domains (IgDs) according to the secondary structure predictions (Figs. 2A and 3A), we performed chemical cross-linking of the TRAPPII complex coupled with mass spectrometry (CXMS) analysis and pull-down experiment to further identify the locations of Trs120 and Trs130. The results of CXMS used by both BS3 and DSS showed cross-links between Trs120 and Trs31, suggesting that Trs120 is close to the Trs31 (fig. S4). Moreover, Trs20 could be pulled down by Trs120-NTS (residue 406-644) indicating the existence of their interaction (fig. S5A). Because the map qualities of the N-terminal region (residues 1-264) are very poor, we could only build the atomic models of the NTS (residues 264-644) and four IgDs (IgD1-4) of Trs120 (Figs. 2, A and B, and fig. S5B). NTS of Trs120 is a slightly curved spiral consisting of about 380 residues arranged into 13 helices (residues 264– 644) and contacts with Trs20 in the TRAPPI (Figs. 1, B and C, and 2B). IgD2-4 contribute to most stable regions of Trs120 (Fig. 2B) and have contacts with both Trs130 and Trs65 (Figs. 1, B and C). Different from the basic structure of IgD, two helices, α1 in IgD2 and α2 in IgD3, inserted into the loop β2-β3 (loop connecting strands β2 and β3) in IgD2 and loop β6-β7 in IgD3, respectively (Fig. 2C). These decorations in loops lead to two long loops, loop α1-β3 of IgD2 and loop α2-β6 of IgD3, stretching toward the interior of the triangle (Fig. 2C). The large gap between these two loops and TRAPPI is supposed to be the position of the bound Ypt31/32, which is confirmed by the resolved structure of the TRAPPII-Ypt32 complex described below.

**Figure 2.**
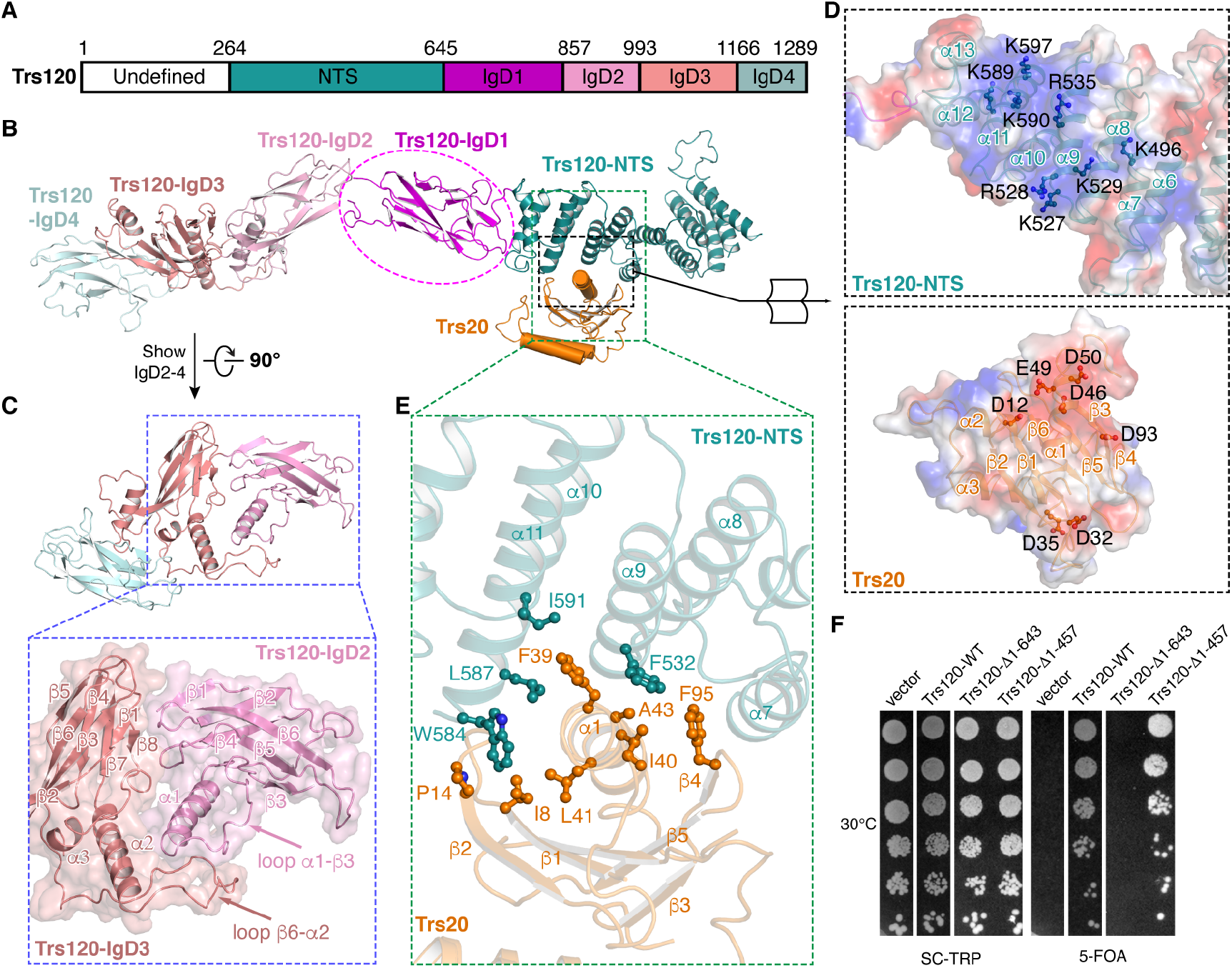
Structure of Trs120 in TRAPPII. **(A)** Schematic representation of the domain structures of Trs120. Color codes for domains are indicated. Numbers indicate the domain boundaries. **(B)** The overall structure of Trs120. **(C)** The interaction between Trs120-IgD2 and Trs120-IgD3. **(D)** Electrostatic surface representation of the interface between Trs120-NTS and Trs20. The surface potentials are complementary. **(E)** The hydrophobic interactions between Trs120-NTS and Trs20. **(F)** Viability of N-terminal deletion mutants of Trs120 tested by yeast survival and growth assays. Cells were grown at 30°C.

**Figure 3.**
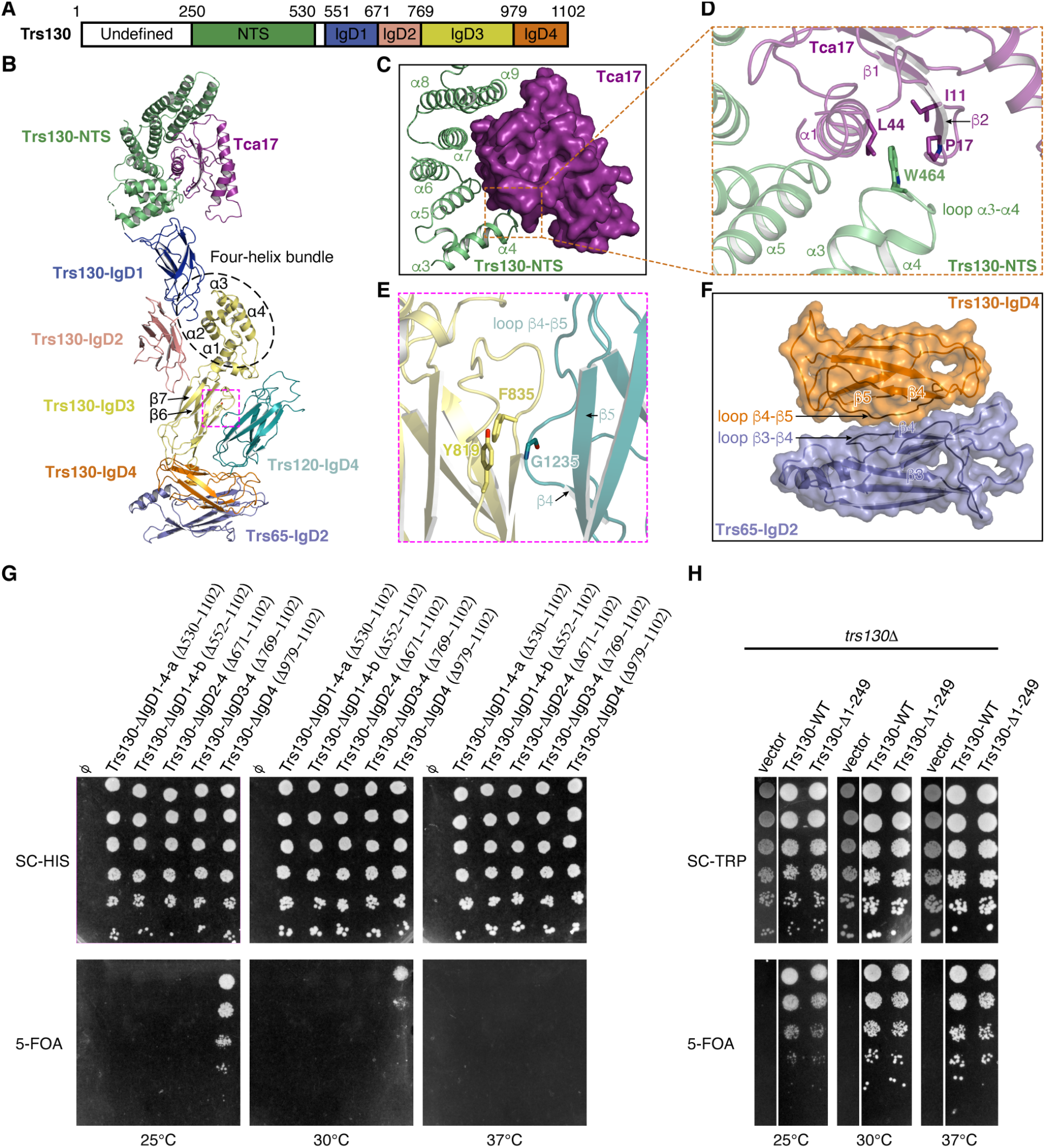
Structure of Trs130 in TRAPPII. **(A)** Schematic representation of the domain structures of Trs130. Color codes for domains are indicated. Numbers indicate the domain boundaries. **(B)** The overall structure of Trs130. The interacting protein Tca17 and protein domains of Trs120-IgD4 and Trs65-IgD2 are also shown. **(C)** The interaction between Trs130-NTS and Tca17. **(D)** W464 of Trs130-NTS is embedded in a hydrophobic pocket formed by residues L44, I11 and P17 of Tca17. **(E)** The interaction between Trs130-IgD3 and Trs120-IgD4. The interaction between the loop β4-β5 of Trs130-IgD4 with the loop β3-β4 of Trs65-IgD2. Viability of C-terminal deletion mutants of Trs130 tested by yeast survival and growth assays. **(H)** Viability of N-terminal deletion mutant of Trs130 tested by yeast survival and growth assays.

Helices α7, α9 and α11 from Trs120-NTS makes extensive interactions with Trs20 (Figs. 2, D and E). In detail, two highly positively charged surfaces of Trs120-NTS contact two acidic patches formed by residues from the helix α1 and loops between strands β1-β2 and between strands β4-β5 of Trs20 (Fig. 2D). Hydrophobic interactions also contribute to the binding between Trs120-NTS and Trs20 as exemplified by the close distances between residues F532, W584, L587 and I591 from Trs120-NTS and I8, P14, F39, I40, L41, A43 and F95 from Trs20 at the interface (Fig. 2E). To further evaluate the physiological relevance of the interaction between Trs120 and Trs20 *in vivo*, we performed growth assays with truncated Trs120. Truncating the N-terminal 457 amino acids of Trs120 that have no interactions with Trs20 slightly affected the yeast growth at various temperatures (Fig. 2F and fig. S5C). Further deleting the N-terminal 643 amino acids that cover all residues interacting with Trs20 caused yeast death at all temperatures (Fig. 2F and fig. S5C). These results suggested that the interactions between Trs120 and Trs20 are essential for yeast survival, probably by maintaining the proper function of TRAPPII.

### Structure of Trs130 in TRAPPII

CXMS analysis also identified seven cross-links between Trs130 and Tca17 (fig. S4), indicating that the super-helical structure near Tca17 is Trs130. According to the density map at this location, we built the almost complete atomic model of Trs130 except for the N-terminal region (residues 1-250) that exhibited no EM density (Figs. 3, A and B, and fig. S5D). The NTS of Trs130 consisting of about 280 residues arranged into 11 helices (residues 250–529) is a more curved spiral than Trs120-NTS (Figs. 2B and 3B). The four IgD domains are connected sequentially and have similar structures except IgD3. IgD3 is distinguished from others by the insertion of a four-helix bundle into the loop β6-β7 (Fig. 3B). This bundle further bridges IgD1 to IgD3 to make the whole structure of IgD1-3 stable (Fig. 3B).

Trs130 makes wide contacts with the surrounding proteins. First, similar with the interaction between Trs120-NTS and Trs20, Trs130-NTS contacts with Tca17 (Figs. 1, B and C, and 3B). Helix α1 of Tca17 is wrapped by a barrel formed by helices α5, α7 and α9 of Trs130-NTS and all four β strands of Tca17 (Figs. 3, B and C). In addition, W464 of Trs130-NTS is embedded in a hydrophobic pocket formed by residues L44, I11 and P17 of Tca17, which is similar with the interaction of W584 of Trs120-NTS with L41, I8 and P14 of Trs20 (Figs. 2E and 3D). Second, IgD3 and IgD4 make up a L-shaped structure holding IgD4 of Trs120 (Fig. 3B). Specifically, the loop between strands β4 and β5 of Trs120-IgD4 extended into the hydrophobic interior (Y819 and F835) between the two β sheets of Trs130-IgD3 (Fig. 3E). Third, the loop between strands β4 and β5 of Trs130-IgD4 contacts with the loop between strands β3 and β4 of Trs65-IgD2 (Fig. 3F). Deletion of Trs130-IgD4 in yeast led to temperature sensitivity and further truncating IgD3 resulted in the yeast death, suggesting that IgD3 was more important to the function of TRAPPII (Fig. 3G). Truncating the N-terminal 249 amino acids of Trs130 affected the growth of yeast slightly in various temperatures (Fig. 3H), which suggested that this unsolved part also has some effects on the function of TRAPPII.

### Structure of Trs65 in TRAPPII

Trs65 composed of three IgDs according to the secondary structure prediction (fig. S6A). CXMS analysis indicated extensive interactions between Trs65 and Trs120 as well as Trs130 (fig. S4). Negative stained images of TRAPPII containing His-GFP tagged Trs65 which is labeled by anti-His antibody showed the Y-shaped antibodies flanked outside TRAPPII, suggesting that the N-terminal IgD1 of Trs65 was located on the lateral side of the TRAPPII complex (fig. S6B). However, its density was too poor to build the atomic model (fig. S6C), likely reflecting its dynamic location due to the long linker between IgD1 and IgD2. Thus, only atomic models of IgD2 and IgD3 of Trs65 were built (fig. S6C). Detailed analysis of the interactions between Trs65 with surrounding proteins suggests that Trs65 could play a critical role in the dimeric complex formation because it interacts with subunits in both monomers (fig. S6D). In one monomer, besides the interaction between Trs65-IgD2 and Trs130-IgD4 as described above (Fig. 3F and fig. S6D), Trs65-IgD3 also contacts with both IgD3 and IgD4 of Trs120. The contact between Trs65-IgD3 and Trs120-IgD3 is mainly mediated by hydrophobic residues (figs. S6, D and E). In addition, helix α1 of Trs65-IgD3 binds to a groove formed by strands β3, β6 and β7, and loop β7-β8 of Trs120-IgD4, in which Y487 of Trs65-IgD3 forms π-π interactions with both F1216 and F1279 of Trs120-IgD4 (figs. S6, D and F). Besides these interactions within the same monomer, Trs65-IgD3 is also very close to the IgD2’ and IgD3’ of Trs120’ in the neighboring monomer suggesting the existence of interactions between them (figs. S6, D and G). Meanwhile, Trs65-IgD2 contacts with Trs120-IgD2’ via the interactions between the loops β3-β4 of Trs65-IgD2 and β2-β1 of Trs120-IgD2’ (figs. S6, D and G). Consistent with the extensive interactions between Trs65-IgD3 and subunits in both monomers, deletion of IgD3 from the yeast genome broke the dimer formation, resulting in a very low abundance of dimers (fig. S6H).

### Structure of the TRAPPII-Ypt32 complex

To further explore how TRAPPII acts as a GEF to activate Ypt32, we resolved the structure of the Ypt32-bound TRAPPII complex (figs. S7, A to D). Similar with the intact TRAPPII complex, the TRAPPII-Ypt32 complex is also a dimer assembled by two triangle shaped monomers (fig. S7F). But different from the Ypt32-free TRAPPII complex in which two distinct conformations of monomer were identified, only one stable closed conformation of monomer was observed in the Ypt32-bound TRAPPII complex (fig. S7B). The monomer was resolved at the resolution of 3.86 Å, which leads to the accurate assignment of all subunits of TRAPPII and the protein of Ypt32 (figs. S7, B to E). The subunit arrangement of TRAPPII in the TRAPPII-Ypt32 complex is almost identical to that of TRAPPII alone (Fig. 4A). As expected, Ypt32 locates inside the triangle-shaped TRAPPII and at the gap between TRAPPI and Trs120 (Fig. 4A). Both the nucleotide-binding domain (NBD) and the C-terminal region of Ypt32 contribute interactions with TRAPPII (Fig. 4A).

**Figure 4.**
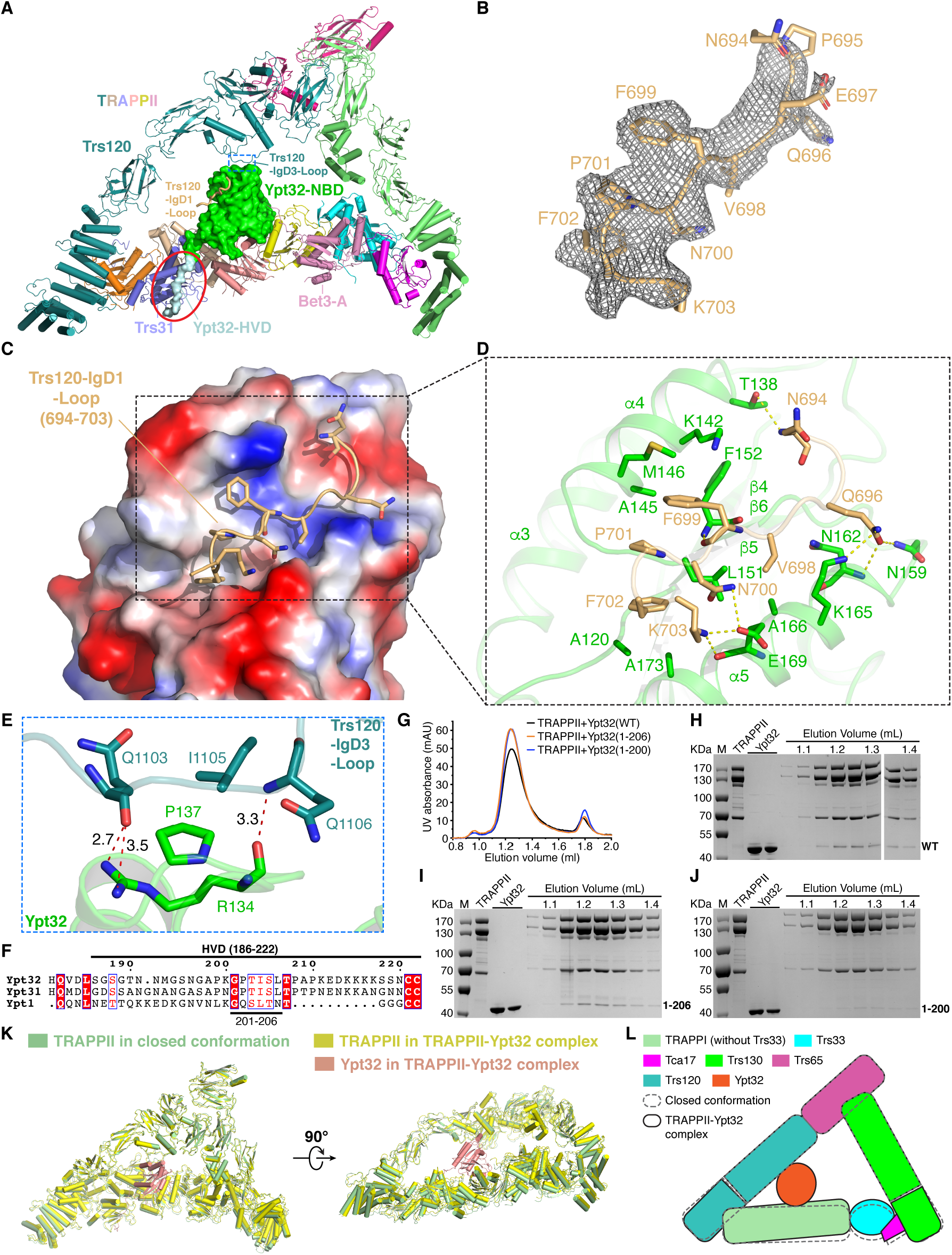
Structure of the TRAPPII-Ypt32 complex. **(A)** The overall structure of the TRAPPII-Ypt32 monomer. TRAPPII is shown in cartoon representation and Ypt32 is displayed in surface representation. The red circle indicates the interaction between the C-terminal of Ypt32 (186-200) and Trs31. **(B)** Cryo-EM densities (mesh) for Trs120-IgD1-Loop. **(C)** Trs120-IgD1-Loop is embedded in one groove of Ypt32. Ypt32 is displayed in the electrostatic surface representation. **(D)** The interactions between Trs120-IgD1-Loop and Ypt32. **(E)** The interactions between Trs120-IgD3-Loop and Ypt32. **(F)** Sequence alignment of C-terminal of Ypt32, Ypt31 and Ypt1 from *Saccharomyces cerevisiae*. **(G)** FPLC curves of the mixtures of TRAPPII incubated with indicated C-terminal deletion mutants of Ypt32. **(H to J)** Peak fractions of the FPLCs were analyzed by SDS-PAGE. **(K)** Structural comparison of the Ypt32-bound TRAPPII and the closed conformation of the Ypt32-free TRAPPII. **(L)** Schematic diagram showing the conformational change between the Ypt32-bound TRAPPII and the closed conformation of the Ypt32-free TRAPPII.

Interactions between NBD of Ypt32 and TRAPPII: There are three contact sites between Ypt32’s NBD and TRAPPII (Fig. 4A). One binding site lies at the interface between NBD of Ypt32 and the TRAPPI core of TRAPPII (Fig. 4A), which is very similar with the interaction between TRAPPI and Ypt1(*22*). The most striking observation is that two additional separated binding sites are formed between NBD of Ypt32 and the TRAPPII-specific component Trs120. We name them TRAPPII-specific binding sites 1 and 2, respectively.

For the TRAPPII-specific binding site 1, the loop (residues N694 to K703) between strands β1 and β2 of Trs120-IgD1, namely Trs120-IgD1-Loop, exactly embeds in the groove formed by strand β5 and helices α4 and α5 of Ypt32 through both hydrophobic and polar interactions (Figs. 4, B to D). The C-terminal part of the Trs120-IgD1-Loop including 4 hydrophobic residues (V698, F699, P701 and F702) is nestled in a hydrophobic surface constituted by A120, A145, M146, L151, F152, A166 and A173 of Ypt32 (Figs. 4, C and D). Two hydrogen bond (H-bond) networks between Trs120-IgD1-Loop and helix α5 of Ypt32 are observed including the interaction of Q696(Trs120)-N159(Ypt32)-N162(Ypt32)-K165(Ypt32), and N700(Trs120)-K703(Trs120)-E169(Ypt32) (Fig. 4D). In addition, the main chain amide group of N694 and carbonyl oxygen (C=O) of F699 from Trs120-IgD1-Loop make H-bonds with the side group of T138 and the main chain amide group of F152 from Ypt32, respectively (Fig. 4D). Moreover, the positively charged Ypt32-K142 forms one cation-π interaction with the aromatic group of Trs120-F699 (Fig. 4D). Sequence alignment indicates that the hydrophobic residues involved in the interaction between Trs120-IgD1-Loop and Ypt32 are more conserved than the charged and polar residues, suggesting that the hydrophobic contacts may play the dominant role in this interaction (figs. S8, A and B).

For the TRAPPII-specific binding site 2, a long loop between helix α2 and strand β6 of Trs120-IgD3 (Trs120-IgD3-Loop), which stretches toward the interior of the triangle-shaped TRAPPII, also has interactions with Ypt32 (Figs. 4, A and E). Arg134 in Ypt32 contributes two H-bonds at this interface. One is formed by the guanidinium group of Arg134 with the main chain C=O of Gln1103 in Trs120, and the other one is between the main chain C=O of Arg134 and the main chain amide groups of Gln1106 in Trs120. In addition, potential hydrophobic interaction occurs between Pro137 in Ypt32 and Ile1105 in Trs120 (Fig. 4E).

Interactions between C-terminal region of Ypt32 and TRAPPII: In our TRAPPII-Ypt32 map, the density for the C-terminal region including the hypervariable domain (HVD) of Ypt32 could be observed, although only partial sequence (186-200) was modeled with polyalanine due to the low resolution. Importantly, the structure clearly shows that this C-terminal region of Ypt32 extends as a rope to attach to the surface of Trs31 (Figs. 4, A and F). CXMS analysis also indicated the cross-linked residue pairs between Lys211 of Ypt32 and Lys168 of Trs31 (fig. S7G). To further investigate the interaction between HVD and TRAPPII, we constructed several C-terminally truncated Ypt32 mutants and tested their binding abilities with TRAPPII (Figs. 4, G to J, and figs. S8, C to E). Deleting the last 10 (Ypt32 1-212) or 16 (Ypt32 1-206) residues at the C-terminal of Ypt32, the protein could still bind to the TRAPPII, and the binding ability was similar to that of wild-type Ypt32 (Figs. 4, H and I, and fig. S8D). However, further deleting 6 more residues (Ypt32 1-200) led to a significant decrease of the binding ability (Fig. 4J). After deleting the whole HVD (Ypt32 1-187), the protein completely lost its ability to bind with TRAPPII (fig. S8E). These results indicate that HVD plays an essential role in the binding of Ypt32 with TRAPPII, and the conserved residues 201-206 of HVD are critical to this interaction.

Two conformations of monomer, open and closed, in the Ypt32-free TRAPPII complex, were observed (Figs. 1, B and C). When we examined the conformational change induced by the binding of Ypt32 to TRAPPII by superimposing TRAPPII-Ypt32 with the open and closed conformations, the results indicate that the Ypt32-bound TRAPPII exhibits the closed conformation (Fig. 4K). Moreover, binding with Ypt32 brings TRAPPI and Trs120 closer (Figs. 4, K and L), since Ypt32 contacts with both of them (Fig. 4A). Therefore, TRAPPI moved slightly toward Trs120, leading to a more compact conformation of TRAPPII-Ypt32 (Figs. 4, K and L).

### Mechanism of Ypt32 activation

To investigate the role of the interactions of the TRAPPII-specific loops with Ypt32, we compared the structures of Ypt32 in the TRAPPII-Ypt32 complex, in the GDP-bound state (PDB:3RWO) and in the GppNHp (a nonhydrolyzable analog of GTP)-bound state (PDB:3RWM) by superposing them (Fig. 5A). As was shown in Fig. 5B, binding with Trs120-IgD1-Loop causes the movement of helices α4 and α5 of Ypt32. Because helix α5 is connected to the SAL motif (residues 155-160), its shift makes the shift of SAL motif. In addition, the movement of helix α4 changes the position of the loop α4-β5, which is also affected by binding with Trs120-IgD3-Loop (Fig. 5B). As the loop α4-β5 interacts with the SAL motif (Fig. 5B), its change also leads to the change of SAL. Thus, all of these drive the relocation of the SAL motif (Fig. 5B). In detail, Ala157 moves toward to the nucleotide-binding pocket, leading to an overclose distance between Ala157 and the guanosine base, which could interfere with the nucleotide binding (Fig. 5B). Moreover, L158, which stabilizes the guanosine base via the hydrophobic contact in the GDP-bound state, moves away from and no longer interacts with the guanosine base (Fig. 5B). Together, binding with Trs120 loops induces the conformational change of the SAL motif of Ypt32 to be ready for the nucleotide release.

**Figure 5.**
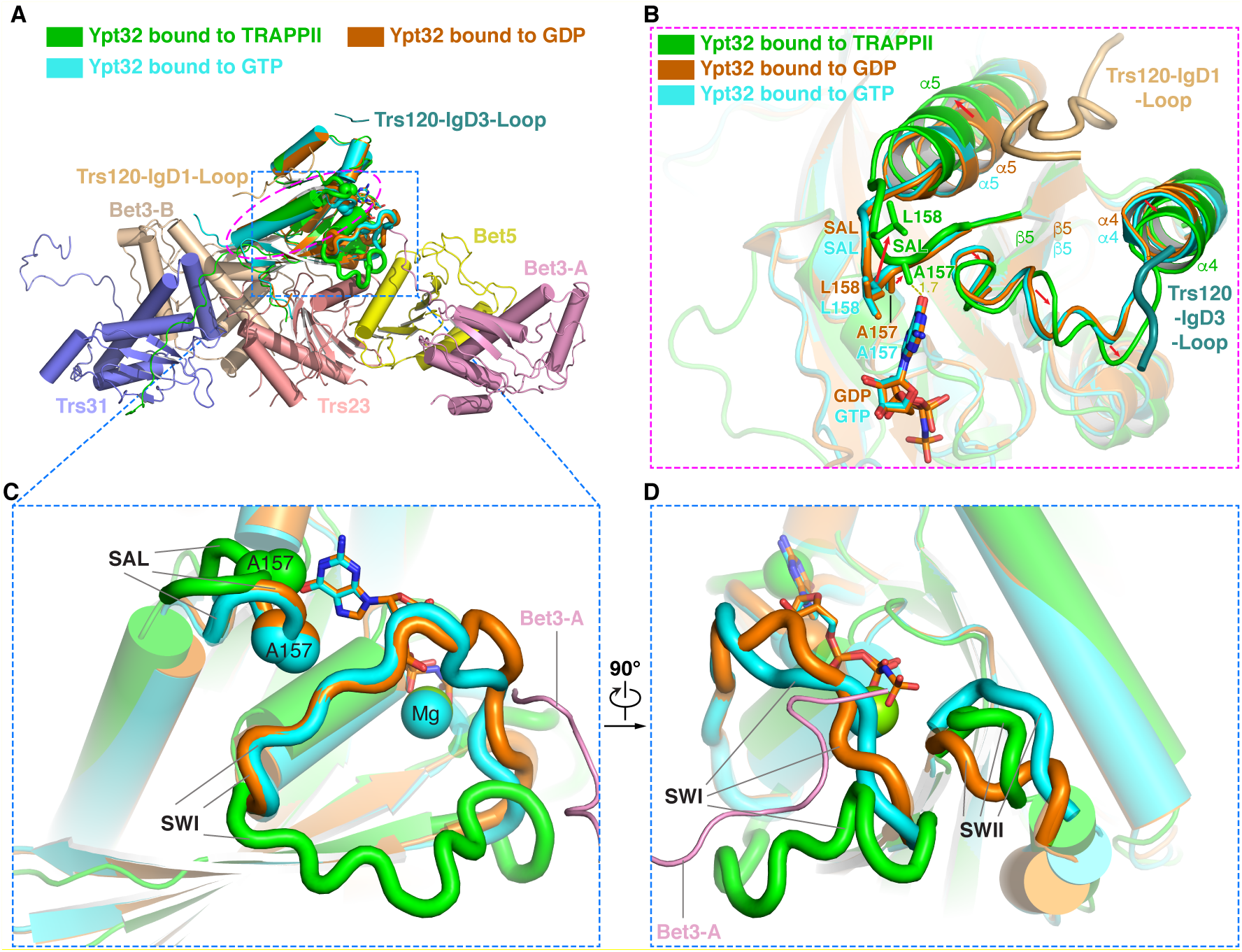
Mechanism of Ypt32 activation. **(A)** Structural comparison of Ypt32 bound to TRAPPII, to GDP (PDB:3RWO) and to GTP (GppNHp, PDB:3RWM). **(B)** Binding with Trs120 loops causes the movement of helices α4 and α5, and loop α4-5 of Ypt32, and the relocation of residues in the SAL motif (red arrows). **(C)** The different positions of SAL motif and SWI motif. **(D)** The different positions of SWI motif and SWII motif.

Except for the change of the SAL motif mentioned above, notable differences also occur in Switch I and II motifs (Figs. 5, C and D). On the Switch I motif (residues 34-48, SWI), due to the steric clash between the C-terminus of Bet3-A and the SWI of the nucleotide-binding Ypt32, SWI of the TRAPPII-bound Ypt32 moves away from the nucleotide-binding pocket, which opens the nucleotide-binding pocket in a favorable conformation for nucleotide release (Figs. 5, C and D). Additional difference occurs on the Switch II motif (residues 68-86, SWII), which locates right between the positions of SWII in GTP-bound and GDP-bound states (Fig. 5D). All of these structural features indicate that the Ypt32 in the TRAPPII-Ypt32 complex is in open conformation, which is consistent with the fact that no nucleotide density was observed in the map of the TRAPPII-Ypt32 complex, which likely represents the intermediate state of GTPase.

## Discussion

In our structure of the TRAPPII-Ypt32 complex, Ypt32 clearly makes contacts with TRAPPI in TRAPPII with the similar mode between Ypt1 and TRAPPI. However, previous studies have shown that TRAPPI alone could only activate Ypt1, but not Ypt32 both *in vitro* and *in vivo* (*11, 22, 24*). To uncover the possible reason behind the different GEF activity between TRAPPI for Ypt1 and TRAPPII for Ypt32, we analyzed several related structures. In the cytosol, the inactive Ypt GTPase is bound with a GDP dissociation inhibitor (GDI) (*27, 28*). Once the GDI-bound Ypt is recruited to the membrane and interacts with GEF, GDP will be released and followed by GTP binding. Thus, we compared the structure of Ypt1 in the GDI-bound state with its structure in the TRAPPI-Ypt1 complex. And we also compared the structure of Ypt31 in the GDI-bound state with the Ypt32’s structure in the TRAPPII-Ypt32 complex, since only the structure of the Ypt31-GDI complex has been resolved and Ypt31 shares over 80% sequence identity with Ypt32 (fig. S9A). We surprisingly found that the loop β2-β3 of Ypt1 moves a lot towards the TRAPPI upon binding to TRAPPI (fig. S9B), whereas this loop of Ypt32 doesn’t change at all upon binding to TRAPPII (fig. S9C). In Ypt1, such movement drives the movement of helix α5 via the polar interaction between D53 in the loop β2-β3 and K170 in the helix α5, which further causes the relocation of the SAL motif to accommodate the nucleotide release (fig. S9B). Different from Ypt1, the loop of Ypt32 remains the same upon binding to TRAPPI in TRAPPII (fig. S9C), and thus could not induce the shift of helix α5 and the change of SAL. However, Ypt32’s SAL in TRAPPII-Ypt32 does show the similar conformation to Ypt1’s SAL in TRAPPI-Ypt1 (fig. S9C). Actually, Trs120, the TRAPPII-specific subunit, plays the role of the relocation of Ypt32’s SAL. As was shown in Fig. 5B, binding with Trs120-IgD1-Loop causes the conformational change of Ypt32’s SAL to adapt a favorable microenvironment for the nucleotide release. Thus, these results may explain why TRAPPI alone is incapable of activating Ypt32, and reveal the critical role of Trs120 in TRAPPII’s activation of Ypt32.

The different conformations of Ypt32-free TRAPPII and Ypt32-bound TRAPPII lead to a possible model of TRAPPII-mediated Ypt32 activating (fig. S9D). First, TRAPPII is recruited onto the trans-Golgi network by anion charge and other factors, such as its regulatory GTPase Arf1 (*24*). As the substrate, the inactive Ypt32 in the cytosol is bound to GDI (*27, 28*). GDI interacts with both NBD and C-terminal hydrophobic prenyl groups of Rab to act as a chaperone protecting the prenylated C terminus and re-extract Rab from membrane (*29*). According to our structure, both NBD and HVD of Ypt32 bind to TRAPPII, but the interface between NBD and TRAPPII is much larger than that between HVD and TRAPPII (Fig. 4A). Therefore, as the second step, the recruitment of Ypt32 to TRAPPII might be initiated by the interaction between NBD of Ypt32 with TRAPPII, which could further induce the dissociation of HVD from GDI. Although two conformations exist in the Ypt32-free TRAPPII, the space of the interior of TRAPPII in the open conformation is larger than that in the closed conformation and the binding site for Ypt32 is more exposed in the open conformation, therefore the open conformation may more readily accommodate the Ypt32 than the closed conformation (Figs. 1, B to D). However, the binding of Ypt32 to TRAPPII in the open conformation is not stable, exemplified by only closed conformation observed in the TRAPPII-Ypt32 complex (Fig. 4K and fig. S7B). Indeed, the dynamic natures of TRAPPII and the binding with Ypt32 provide the opportunity for the Ytp32-bound TRAPPII in the open conformation to switch to the closed conformation. Along with this transition, the TRAPPII moves closer to the membrane and the bound Ypt32 is activated for the nucleotide exchange (*30*).

## Acknowledgments

We are grateful to J. Wang for model building. We thank K. Mei for helpful discussion. We thank the staff at the Tsinghua University Branch of the National Protein Science Facility (Beijing) for their technical support on the Cryo-EM and High-Performance Computation platforms.

## Author contributions

S.-F.S. supervised the project; C.M., L.Z. and G.H. prepared the samples, performed the biochemical and yeast growth analyses; C.M. and F.Y. collected the EM data and performed the EM analysis; C.M., L.Z. and X.Y. performed the model building and the structure refinement; G.S. and M.-Q.D. designed and performed the CXMS assays; C.M., S.S. and S.-F.S. analyzed the structure; C.M. wrote the initial draft; S.S and S.-F.S. edited the manuscript.

## Data and materials availability

The EM density map of the intact dimeric TRAPPII complex has been deposited in the Electron Microscopy Data Bank (www.ebi.ac.uk/pdbe/emdb/). Atomic coordinates and EM density maps of the intact dimeric TRAPPII complex in State I (PDB: XXXX; EMDB: EMD-XXXX), in State II (PDB: XXXX; EMDB: EMD-XXXX), in State III (PDB: XXXX; EMDB: EMD-XXXX), the monomer of TRAPPII in the closed conformation (PDB: XXXX; EMDB: EMD-XXXX), in the open conformation (PDB: XXXX; EMDB: EMD-XXXX), the intact dimeric TRAPPII-Ypt32 (PDB: XXXX; EMDB: EMD-XXXX) and the monomer of TRAPPII-Ypt32 (PDB: XXXX; EMDB: EMD-XXXX) have been deposited in the Protein Data Bank (www.rcsb.org) and the Electron Microscopy Data Bank (www.ebi.ac.uk/pdbe/emdb/). All other data and materials are available from the corresponding authors upon reasonable request.

## Supplementary Materials

Materials and Methods

Figures S1-S9

Tables S1-S2

