## Supplementary Information for "Structural basis for assembly of TRAPPII complex and specific activation of GTPase Ypt31/32"

**This PDF file includes:**

Materials and Methods
Figs. S1 to S7
Tables S1 and S2

### Materials and Methods

Clones and plasmids

Strains with C-terminal Flag-tagged Trs120 or with gene sequence deletion or truncation were constructed by lithium acetate method (*31*). The genome of *S. cerevisiae* cells was as a PCR template for all plasmid construction. Gene sequence of Trs20 and residues 406-644 of Trs120 were optimized for exogenous expression of *Escherichia coli.*

Purification of TRAPPII

The Trs120-3xFLAG-tagged *S. cerevisiae* (44 liters) were grown in YPD medium at 30℃ overnight until an $OD_{600}$ reached to 5~6. The cells were pelleted by centrifugation and resuspended using the lysis buffer (20 mM HEPES-NaOH pH 7.4, 150 mM NaCl, 10% glycerol, 1% Chaps, 1 mM EDTA, 2 mM DTT, 1 mM PMSF) supplemented with cocktail (Roche). The cell pellets were frozen by liquid nitrogen, then lysed by Freezer Mill 6875 (SPEX CertiPrep). The lysate was centrifuged at 35000 rpm for 40 min. All supernatants were incubated with anti-Flag affinity resin (Sigma) at 4℃ for 2 h. The resin was then washed by the washing buffer (20 mM HEPES-NaOH pH 7.4, 300 mM NaCl, 10% glycerol, 0.1% Chaps, 1 mM EDTA, 2 mM DTT) and eluted with the washing buffer supplemented with 0.1 mg/ml Flag peptide (Sigma). The eluate was concentrated and applied to a Superose 6 Increase 3.2/300 (GE Healthcare) equilibrated with the sample buffer (20 mM HEPES-NaOH pH 7.4, 150 mM NaCl, 2 mM DTT, 0.05% digitonin). The peak fractions were analyzed by SDS-PAGE and the mass spectrometry (MS) analysis.

Preparation of TRAPPII-Ypt32 complex

*Escherichia coli* BL21 (DE3) transformed by His-sumo-Ypt32 plasmid were grown at 37℃, then induced by 0.5 mM IPTG at 16℃ overnight and purified by Ni-beads (GE Healthcare). After binding with beads, the beads were washed by the basic buffer (20 mM HEPES-NaOH pH 7.4, 300 mM NaCl, 5% glycerol) with 20mM imidazole and eluted with the basic buffer supplemented with 300mM imidazole (Sigma). The elute was then applied to a Superdex 200 Increase 3.2/300 (GE Healthcare) equilibrated with the incubation buffer (20 mM HEPES-NaOH pH 7.4, 150 mM NaCl, 5% glycerol, 2 mM DTT) with 20 mM EDTA and 0.05% digitonin. TRAPPII complex was changed into the same buffer. To prepare the TRAPPII-Ypt32 complex, TRAPPII complex and his-sumo-Ypt32 were incubated at 4℃ for one and a half hours, and then loaded onto a Superose 6 Increase 3.2/300 (GE Healthcare) equilibrated with the elution buffer (20 mM HEPES-NaOH pH 7.4, 150 mM NaCl, 2 mM DTT) with 0.05% digitonin. Peak fractions were analyzed by SDS-PAGE. His-sumo-Ypt32 with C-terminal truncation in various length were prepared as above and analyzed by SDS-PAGE.

Cryo-EM sample preparation and processing

Aliquots (~4 µl) of TRAPPII complex (~1.5 mg/ml) or TRAPPII-Ypt32 complex (~1.7 mg/ml) were applied to freshly glow-discharged holey carbon grids (Quantifoil Au R1.2/1.3 400 mesh) and blotted for 3.5 s in 100% humidity at 8℃. The grids were plunged into liquid ethane by a Mark IV Vitrobot (FEI). Raw micrographs of TRAPPII complex were collected using a Titan Kiros Microscope (Thermo Fisher Scientific) operated at 300 kV and equipped with a K2 Summit direct electron detector (Gatan) and a GIF Quantum energy filter (Gatan). The cryo-EM images were automatically collected using AutoEMation (*32*) with a slit width of 20 eV on the energy filter and a preset defocus range of −1.8 µm to −1.3 µm in super-resolution mode at a nominal magnification of 105,000×. A total dose of approximately 50 electrons per $Å^{-2}$ for each movie stack was fracted into 32 frames over 5.6 s exposure time. The stacks were motion-corrected with MotionCor2 (*33*) and binned twofold, resulting in a pixel size of 1.091 Å. Raw micrographs of TRAPPII-Ypt32 complex were collected using the same microscope (FEI) equipped with a K3 Summit direct electron detector (Gatan). The cryo-EM images were automatically collected using AutoEMation with a slit width of 20 eV on the energy filter and a preset defocus range of −1.8 μm to −1.3 μm in super-resolution mode at a nominal magnification of 81,000×. A total dose of approximately 50 electrons per Å$^{-2}$ for each movie stack was fracted into 32 frames over 2.56 s exposure time. The stacks were motion-corrected with MotionCor2 and binned twofold, resulting in a pixel size of 0.8697 Å.

Image processing

Nearly all steps of image processing were using RELION (*34-36*). Contrast transfer function (CTF) parameters were estimated by CTFFIND4 (*37*). For TRAPPII complex, we first manually picked and extracted about 2000 particles, which were applied to the reference-free 2D classification. The obtained class averages were used as templates for automatic particle picking in RELION. All auto-picked particles were subjected to several rounds of 2D classification to remove obvious poor particles including ice contaminants, aggregates, and further checked manually, after which a set of 809,002 particles were retained. 2,400 particles from 2D classification were selected to generate an initial model using RELION which was low-pass filtered to 60 Å to be used as the template for 3D classification. After 2 rounds of 3D classification, 178,627 particles were selected and produced a 3D reconstruction with an average resolution of 3.87 Å without imposing any symmetry. A local mask of center region was applied during the refinement and improved the resolution to 3.44 Å (Extended Data Fig. 1e). The density map showed an apparent two-fold symmetry, but only one half displayed clear density and the other half exhibited very poor density suggesting its heterogeneity. To deal with this, we expanded the data set by rotating one monomer (half of the entire dimer) 180° along the C2 symmetry axis by adding 180 to the value of the column _rlnAngleRot in the star file of particles, so that both monomers are reoriented onto a single position. We then performed classification by applying an extraordinarily soft mask around the reoriented monomers. Through this strategy, two distinct conformations of monomer were identified, and 3D refinement yielded the open and closed structures at 4.15 Å and 3.71 Å resolutions, respectively (Extended Data Fig. 1f). We further assigned each monomer back to its original dimeric particle, resulting in the reconstruction of TRAPPII structure in three different states at resolutions of 4.36 Å, 4.67 Å and 6.54 Å for State I, II and III, respectively (Extended Data Fig. 1f). Local resolution was estimated by ResMap(*38*).

For TRAPPII-Ypt32 complex, particles were automatically picked up using the same templates as TRAPPII in RELION. After 2D classification, all particles were checked manually and resulted in a set of 162,668 particles. After 3D classification, 56,658 particles were selected. Using the same expanded-symmetry 3D classification strategy, we obtained one stable conformation of monomer at 3.85 Å (Extended Data Fig. 6b). By assigning each monomer back to its original dimeric particle, the whole structure of TRAPPII-Ypt32 was resolved at 4.46 Å with imposing C2 symmetry (Extended Data Fig. 6b). Local mask of different subunits were applied to improve the quality of map density for model building.

<u>Model building</u>

The density maps of the TRAPPII monomer in different conformations and the TRAPPII-Ypt32 complex were used for the model building. The atomic models were generated by a strategy combining rigid body fitting, homology modelling and *de novo* modelling. Briefly, the crystal structures of TRAPPI subcomplexes (PDB 3CUE, 2J3T and 2J3U) and Tca17(PDB 3PR6) were docked into the map using CHIMERA (*39*), and then the structure were manually rebuilt and adjusted based on the density map in COOT (*40*). Atomic models of Trs65, Trs120 and Trs130 were built *de novo.* The results of MS analysis of cross-linked TRAPPII complex provided hints for identifying those three important components. Secondary structure prediction by Phyre2 (*41*) aided the main-chain tracing. After poly-Ala backbones were built manually, different domains were searched by DALI server (*42*) in order to find ideal homologue models. Sequence assignments were guided mainly by bulky residues such as Trp, Tyr, Phe, Arg and Lys. Models were refined using phenix.real_space_refinement against masked map with applying secondary structure and stereochemical constraints (*43*). In case of possible clashes between different domains, combine_focused_map in Phenix was used (*43*). Finally, most residues were assigned for all subunits with some residues were presented as poly-Ala. Residues 1-211 of Trs65, 1-263 of Trs120, 1-249 of Trs130, and 674-693 and 704-728 of Trs120 could not be modelled due to the poor density maps. For TRAPPII-Ypt32 complex, Ypt32 in GDP/GTP form (PDB: 3RWM and 3RWO) and TRAPPII of closed conformation were docked into the map and manually rebuilt and adjusted in COOT. The final atomic models of TRAPPII in closed conformation and open conformation and the TRAPPII-Ypt32 monomer were cross-validated according to previously described procedures (*44, 45*). Briefly, atoms in the model were randomly shifted by up to 0.5 Å, and then refined against one of the two independent half maps generated during the final 3D reconstruction. Then, the refined model was tested against the other map. To obtain the atomic models of the intact TRAPPII and TRAPPII-Ypt32 complex, atomic models of monomers were docked into the density maps of the dimeric TRAPPII in different states and the dimeric TRAPPII-Ypt32 complex. The data collection, model refinement and validation statistics are presented in Extended Data Table 1. The model building is summarized in Extended Data Table 2. The statistics of the geometries of the models were generated using MolProbity (*46*).

The sequence alignments were performed by Clustal W (*47*) and created by ESPript (*48*). All figures and movies were prepared using CHIMERA or PYMOL (http://www.pymol.org).

Chemical Cross-linking mass spectrometry (CXMS) analysis

For CXMS analysis, about 10 μg of TRAPPII was cross-linked with 1 mM DSS or BS3 for 1 h at 25℃, and about 10 μg of TRAPPII-Ypt32 was cross-linked with 1 mM DSS for 1 h at 25℃. The reactions were quenched with 20 mM $NH_4HCO_3$. Proteins were precipitated with ice-cold acetone, resuspended in 8 M urea, 100 mM Tris pH 8.5, and then digested by trypsin (Promega) in 2 M urea, 100 mM Tris (pH 8.5). The LC-MS/MS analysis was performed on an Easy-nLC 1000 II HPLC (Thermo Fisher Scientific) coupled to a Q-Exactive HF mass spectrometer (Thermo Fisher Scientific). Peptides were loaded on a pre-column (75 μm ID, 6 cm long, packed with ODS-AQ 120 Å–10 μm beads from YMC Co., Ltd.) and further separated on an analytical column (75 μm ID, 13 cm long, packed with Luna C18 1.9 μm 100 Å resin from Welch Materials) with a linear reverse-phase gradient from 100% buffer A (0.1% formic acid in H2O) to 30% buffer B (0.1% formic acid in acetonitrile) in 56 min at a flow rate of 200 nl/min. The top 15 most intense precursor ions from each full scan (resolution 60,000) were isolated for HCD MS2 (resolution 15,000; normalized collision energy 27) with a dynamic exclusion time of 30 s. Precursors with 1+, 2+, 7+ or above, or unassigned charge states were excluded. The pLink (*49*) software was used to identify cross-linked peptides with precursor mass accuracy at 20 ppm, fragment ion mass accuracy at 20 ppm, and the results were filtered by applying a 5% FDR cutoff at the spectral level and then an E-value cutoff at 0.001 (*50*).

GST pull-down assay

Constructions of the N-terminal GST-fused Trs120 (406-644) and the N-terminal His-Sumo tagged Trs20 were co-transformed into *Escherichia coli* BL21 (DE3) cells. When the cells were cultured to an $OD_{600}$ of 0.6~0.8, the protein was induced by adding 0.5 mM IPTG and the cells continued to grow at 16℃ overnight. The proteins were purified using glutathione agarose (GE Healthcare). Transformation of only His-Sumo-Trs20 was used as the control to ensure that His-Sumo-Trs20 cannot interact with glutathione agarose. The bound protein complexes were detected by SDS-PAGE.

Antibody labeling

A Yeast strain in which Trs65 gene was deleted from the genome and contained C-terminal 3xFlag tagged Trs120 was constructed and named Trs65del.

The construction of 6xHIS-GFP-Trs65 drived by GAL1 promotor was transformed into yeast strain Trs65del. 12 L of transformed yeast cells were cultured in minimal medium lacking leucine with supplement of raffinose until the $OD_{600}$ reached to logarithmic growth phase, then galactose was added to the final concentration of 100 mM. After 6 hours of culture, the cells were pelleted and proteins were purified by anti-Flag affinity resins (Sigma). Anti-HIS antibodies (ZSGB-BIO) were added at 1:1.5 molar ratio (antibody to protein) into the purified protein and incubated for 30 min at room temperature. The sample was concentrated and applied to a Superose 6 Increase 3.2/300 (GE Healthcare). The peak fraction was analyzed by negative stain EM.

Dilution plating assays

Yeast strains with N-terminal or C-terminal truncation of Trs120 or Trs130 were produced via plasmid shuffling. Trs120 and Trs130 were separately constructed into P414/P416 plasmid under the control of the ADH promoter and the CYC1 terminator. The chromosomal gene Trs120 and Trs130 were disrupted with a HIS3 cassette. For growth analysis, cells were grown at 30℃, then washed by sterilized water and diluted to $OD_{600}$ of 1.25. The cells were plated at 5-fold serial dilutions on SC-HIS/TRP plates with or without 5-FOA at 25℃, 30℃, 37℃ for 72h. Three independent experiments were performed, and one representative is presented in the figure.

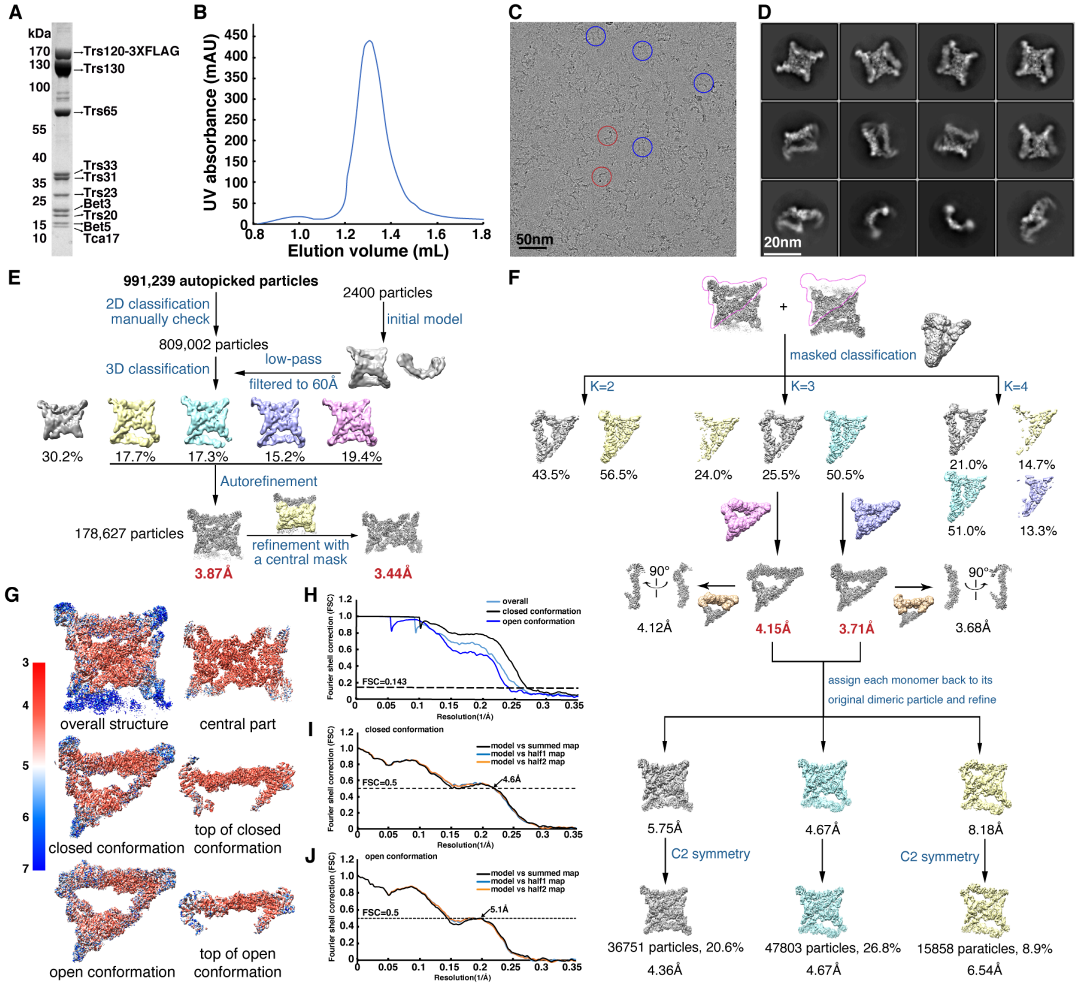

**Fig. S1. Cryo-EM analysis of TRAPPII. (A)** SDS–PAGE analysis of protein components in the purified yeast TRAPPII. The gel was stained with Coomassie brilliant blue. The bands were identified by MS analysis. (**B**) The last-step size exclusion chromatography purification of TRAPPII. (**C**) The representative motion-corrected electron micrograph of TRAPPII with typical particles marked by blue circle (top view) and red circle (side view). (**D**) Typical good reference-free 2D class averages of TRAPPII. (**E**) Typical good reference-free 2D class averages of TRAPPII. (**F**) The flowchart for EM data processing of the monomer of TRAPPII in different conformations. Details can be found in Methods. (**G**) The density maps colored by local resolution. (**H**) Gold-standard Fourier shell correlation (FSC) curves for the 3D electron microscopy reconstructions of the intact dimeric TRAPPII (light blue), the monomer in closed conformation (black) and the monomer in open conformation (blue). **(I to J)** The cross-validations of the atomic models of the TRAPPII monomers in the closed conformation **(I)** and the open conformation **(J)**. FSC curves of the refined model versus the overall map that it was refined against (black), of the model refined against the first half map versus that same map (blue), and of the model refined against the first half map versus the second map (orange). The small difference between the blue and orange curves indicates that the refinement of the atomic coordinates was not affected by overfitting.

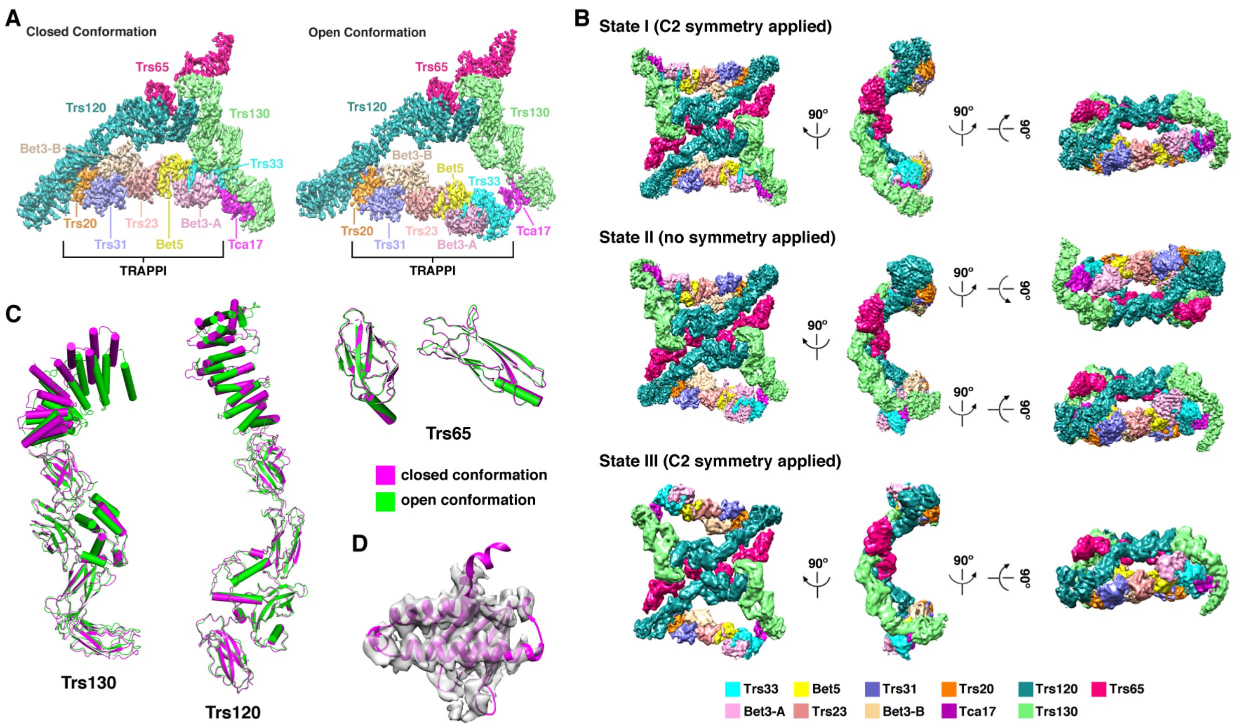

**Fig. S2. 3D reconstruction of TRAPII. (A)** 3D cryo-EM density maps of the TRAPPII monomers in the closed conformation (left panel) and the open conformation (right panel). **(B)** 3D cryo-EM density maps of the intact dimeric TRAPPII in State I (upper panel), State II (middle panel) and State III (lower panel). **(C)** Structural superimposition of Trs130, Trs120 and Trs65 in the closed conformation (magenta) and the open conformation (blue). **(D)** The crystal structure of yeast Tca17 (PDB 3PR6) can be fitted well with the density map.

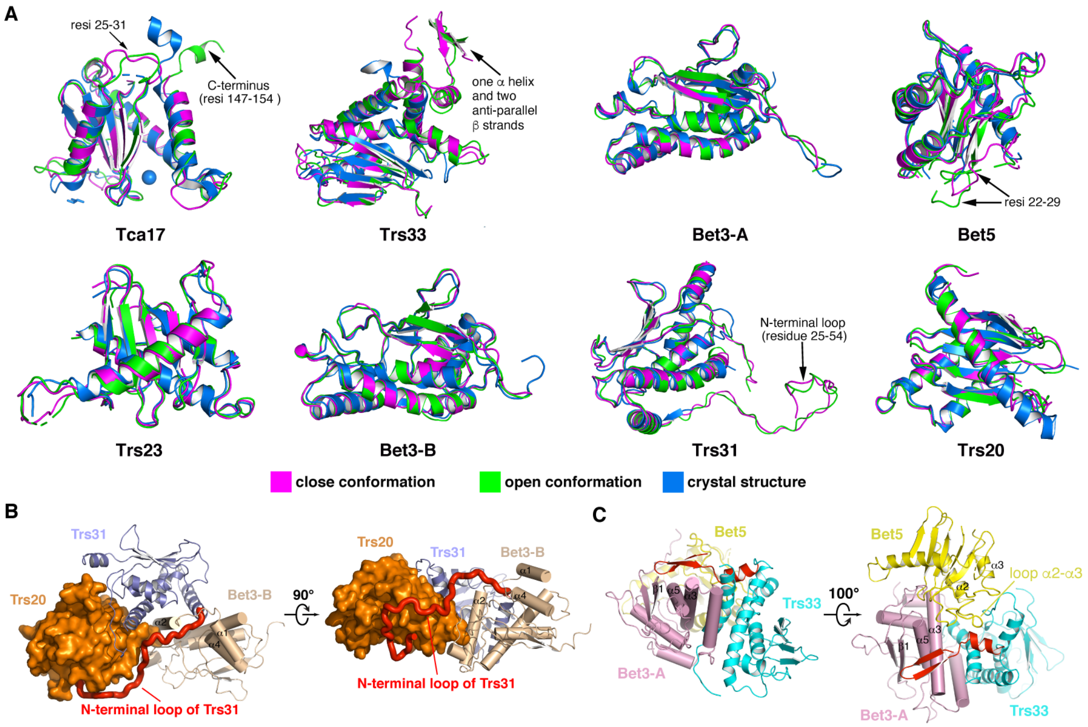

**Fig. S3. Structures of TRAPPI and Tca17 in TRAPPII.** (**A**) Structural comparison of the crystal structures of Tca17 and subunits of TRAPPI with their structures in the different conformations of TRAPPII monomer. (**B**) The interaction between the N-terminal loop of Trs31and Bet3-b and Trs20. (**C**) The extra two β strands and one short α helix (red) of yeast Trs33 interact with Bet3-A and Bet5, respectively.

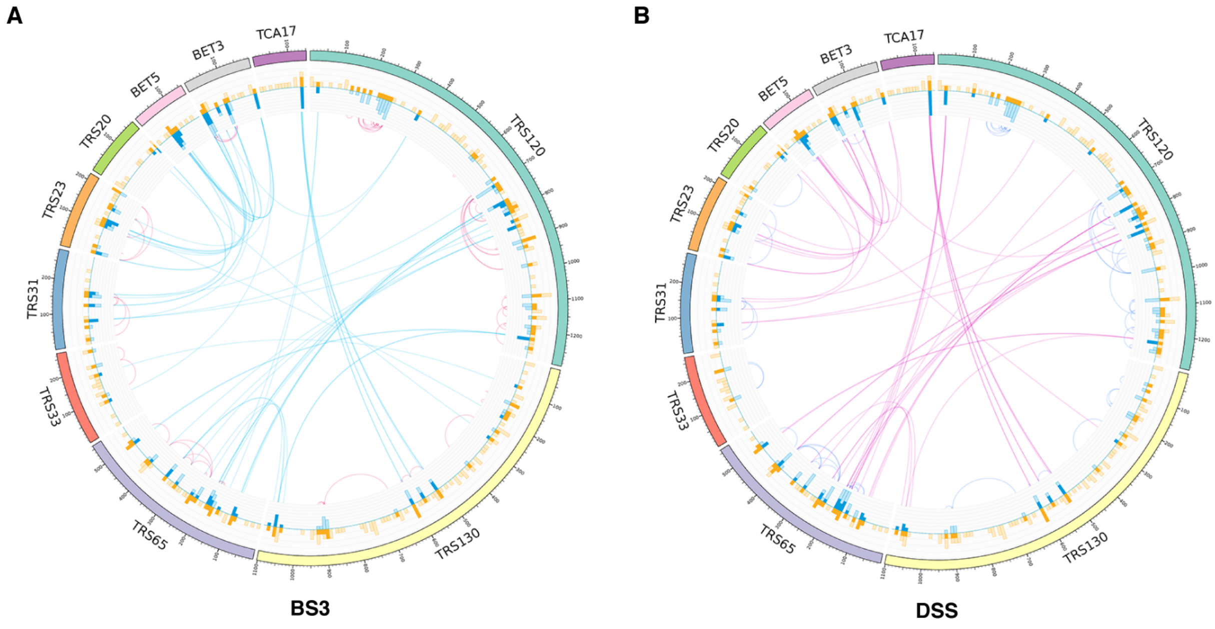

**Fig. S4. Summary of the chemical cross-linking results of TRAPPII.** (**A**) Circular plot showing the distribution of the identified BS3 cross-linked residue pairs mapped to protein sequences. (**B**) Circular plot showing the distribution of the identified DSS cross-linked residue pairs mapped to protein sequences.

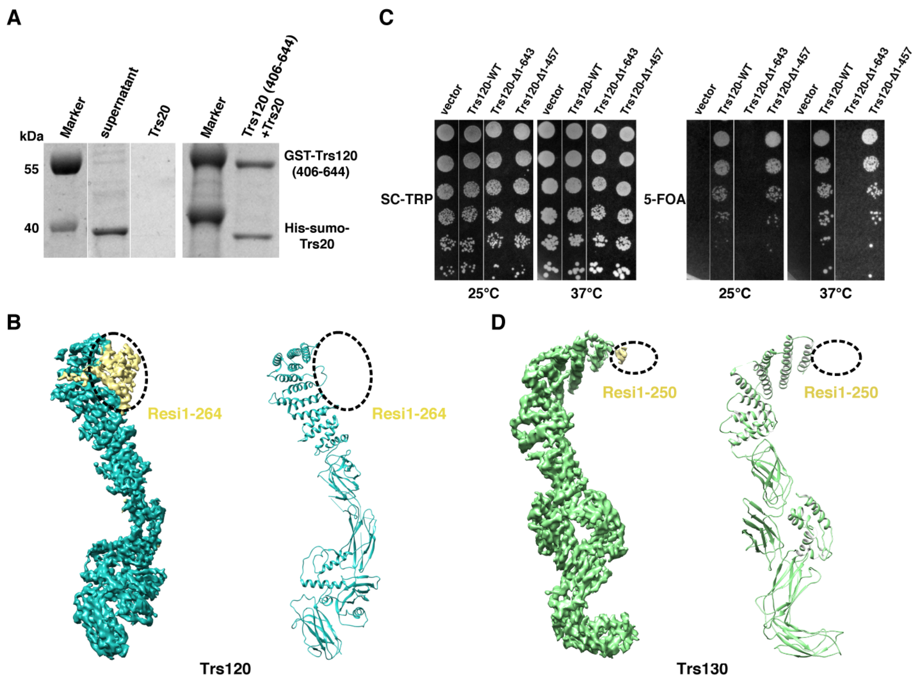

**Fig. S5. Structural features of Trs120, Trs130 and Trs65.** (**A**) Pull-down analysis indicates the interaction between Trs120-NTS and Trs20. (**B**) The density map and atomic model of Trs120. The dashed circles indicate the poor qualities of the density map (left panel, yellow) and the unmodeled sequence (right panel). (**C**) Viability of N-terminal deletion mutants of Trs120 tested by yeast survival and growth assays. Cells were grown at 25℃ or 37℃. (**D**) The density map and atomic model of Trs130. The dashed circles indicate the poor qualities of the density map (left panel, yellow) and the unmodeled sequence (right panel).

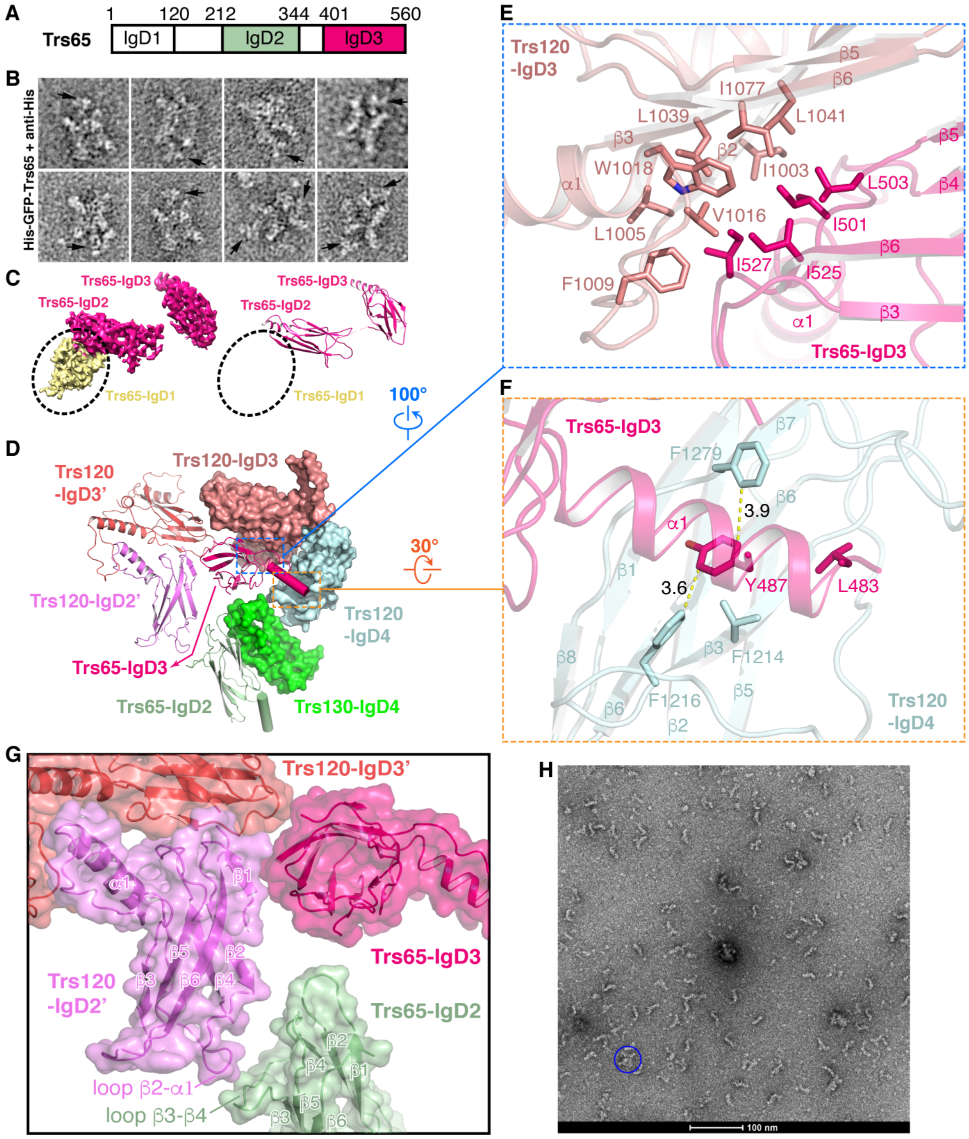

**Fig. S6. Structure of Trs65 in TRAPPII.** **(A)** Schematic representation of the domain structures of Trs65. Color codes for domains are indicated. Numbers indicate the domain boundaries. **(B)** Negative stained images of TRAPPII containing His-GFP tagged Trs65 which is labeled by anti-His antibody. Black arrows indicate the Y-shaped antibodies flanking outside TRAPPII, suggesting that the N-terminal IgD1 of Trs65 was located on the lateral side of the TRAPPII complex. **(C)** The density map and atomic model of Trs65. The dashed circles indicate the poor qualities of the density map (left panel, yellow) and the unmodeled sequence (right panel). **(D)** Overall structures of Trs65 and its interacting proteins. **(E)** The interaction between Trs65-IgD3 and Trs120-IgD3. **(F)** The interaction between Trs65-IgD3 and Trs120-IgD4. **(G)** The interaction between Trs65 with Trs120' in the neighboring monomer. **(H)** Negative stained micrograph of TRAPPII extracted from yeast cells in which the IgD3 of Trs65 was deleted from genome. Most particles are monomers with shapes of triangle or stick. A very low abundance of dimers (blue circle) were observed.

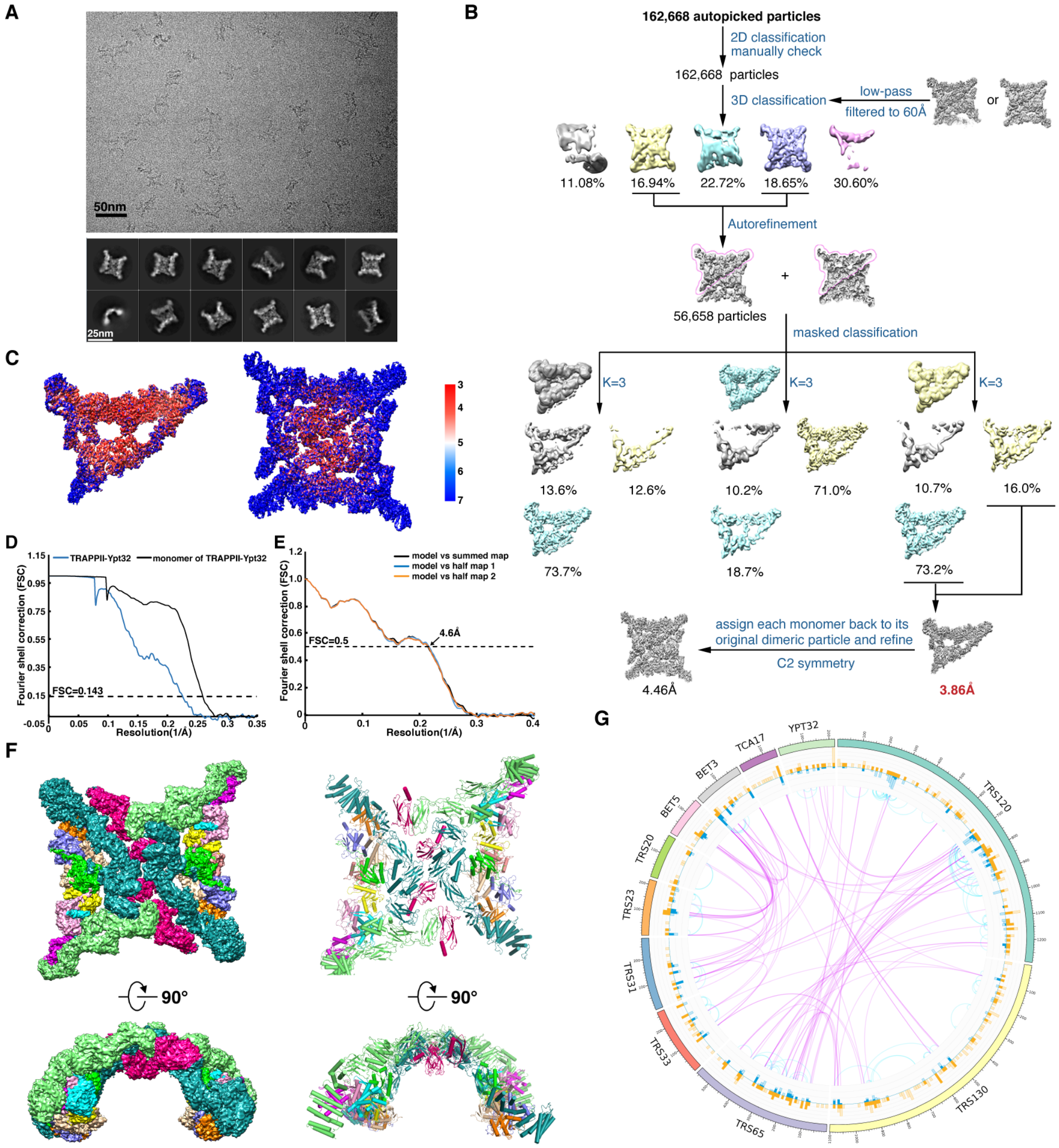

**Fig. S7. Cryo-EM analysis of the TRAPPII-Ypt32 complex. (A)** The representative motion-corrected electron micrograph of TRAPPII-Ypt32 (top panel) and typical good reference-free 2D class averages (lower panel). **(B)** The flowchart for EM data processing of the TRAPPII-Ypt32 complex. **(C)** The density maps colored by local resolution. **(D)** FSC curves for the 3D electron microscopy reconstructions of the intact dimeric TRAPPII-Ypt32 and the monomer. **(E)** The cross-validation of the atomic model of the TRAPPII-Ypt32 monomers. FSC curves of the refined model versus the overall map that it was refined against (black), of the model refined against the first half map versus that same map (blue), and of the model refined against the first half map versus the second map (orange). The small difference between the blue and orange curves indicates that the refinement of the atomic coordinates was not affected by overfitting. **(F)** The density maps (left panel) and atomic models (right panel) of the intact dimeric TRAPPII-Ypt32 complex. **(G)** Summary of the chemical cross-linking results of the TRAPPII-Ypt32 complex. Circular plot showing the distribution of the identified DSS cross-linked residue pairs mapped to protein sequences.

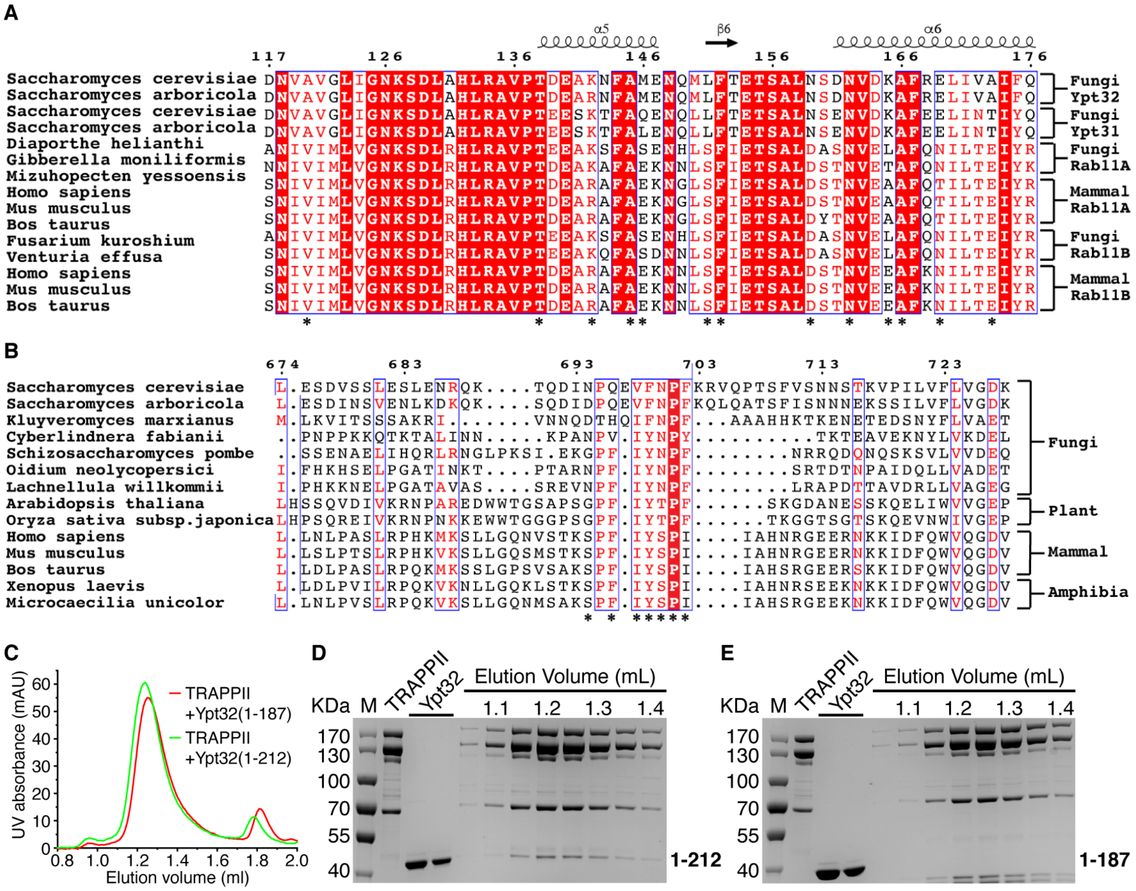

**Fig. S8. The interaction between Trs120 and Ypt32. (A, B)** Sequence alignment of Ypt31/32 **(A)** and Trs120 **(B)** from different species. The residues involved in the interaction are indicated by asterisks. **(C)** FPLC curves of the mixtures of TRAPPII incubated with indicated C-terminal deletion mutants of Ypt32. **(D, E)** Peak fractions of the FPLCs were analyzed by SDS-PAGE.

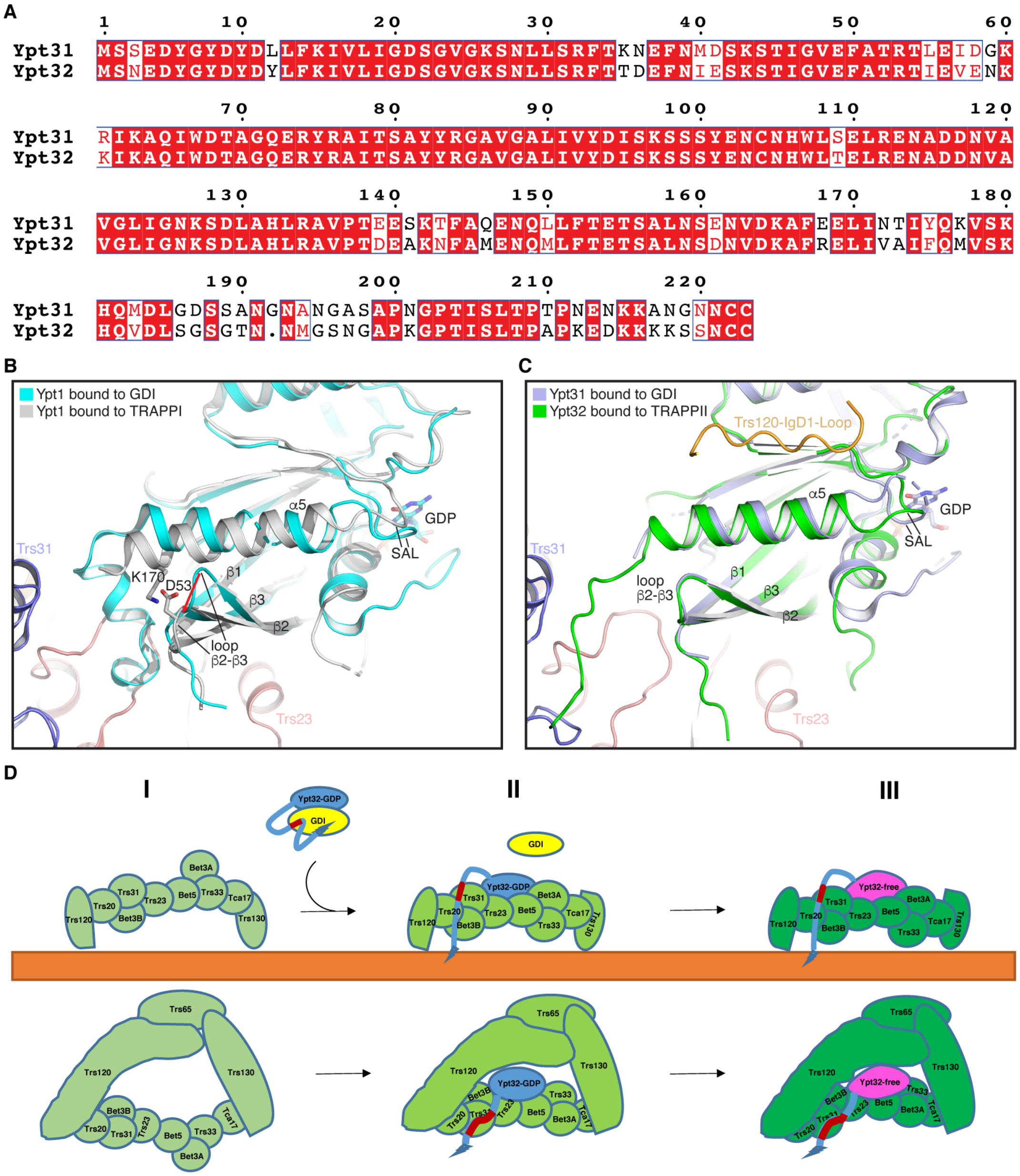

**Fig. S9. Model of TRAPPII-mediated Ypt32 activating. (A)** Sequence alignment of Ypt31 and Ypt32 from yeast. **(B)** Structural comparison of Ypt1 bound to GDI and to TRAPPI. **(C)** Structural comparison of Ypt31 bound to GDI and Ypt32 bound to TRAPPII. **(D)** TRAPPII is recruited onto the trans-Golgi network (I). Then Ypt32/GDI binds to TRAPPII, which could induce the dissociation of HVD from GDI. The released HVD, especially the sequence of G201-L206 that is indispensable for binding of Ypt32 to TRAPPII, could contact to Trs31 in TRAPPII. Along with the transition from the open conformation to the closed conformation, TRAPPII and bound Ypt32 move closer to the membrane, and the shortened distance ensures that the prenylated C-terminus of Ypt32 is inserted into the membrane (II). Also, in the closed conformation, Ypt32 could make additional interaction with Trs120. All these interactions may enable TRAPPII to activate Ypt32 by causing the conformational changes of Ypt32 that allow the nucleotide exchange (III).

**Table S1.**

**Cryo-EM data collection, refinement and validation statistics**

| | Monomer of TRAPPII (closed) (EMDB-XXXX) (PDB XXXX) | Monomer of TRAPPII (open) (EMDB-XXXX) (PDB XXXX) | Intact TRAPPII (dimer) (EMDB-XXXX) (PDB XXXX) | Intact TRAPPII (State I) (EMDB-XXXX) (PDB XXXX) | Intact TRAPPII (State II) (EMDB-XXXX) (PDB XXXX) | Intact TRAPPII (State III) (EMDB-XXXX) (PDB XXXX) | Monomer of Ypt32-TRAPPII (EMDB-XXXX) (PDB XXXX) | Intact Ypt32-TRAPPII (dimer) (EMDB-XXXX) (PDB XXXX) |
|---|---|---|---|---|---|---|---|---|
| **Data collection and processing** | | | | | | | | |
| Magnification | 105,000 | 105,000 | 105,000 | 105,000 | 105,000 | 105,000 | 81,000 | 81,000 |
| Voltage (kV) | 300 | 300 | 300 | 300 | 300 | 300 | 300 | 300 |
| Electron exposure (e-/Å$^2$) | 50 | 50 | 50 | 50 | 50 | 50 | 50 | 50 |
| Defocus range (μm) | 1.3-1.8 | 1.3-1.8 | 1.3-1.8 | 1.3-1.8 | 1.3-1.8 | 1.3-1.8 | 1.3-1.8 | 1.3-1.8 |
| Pixel size (Å) | 1.091 | 1.091 | 1.091 | 1.091 | 1.091 | 1.091 | 0.8697 | 0.8697 |
| Symmetry imposed | C1 | C1 | C1 | C2 | C1 | C2 | C1 | C2 |
| Initial particle images | 809,002 | 809,002 | 809,002 | 809,002 | 809,002 | 809,002 | 158,626 | 158,626 |
| Final particle images | 181,003 | 91,346 | 178,627 | 36,751 | 47,803 | 15,858 | 81,870 | 32,394 |
| Map resolution (Å) | 3.71 | 4.15 | 3.87 | 4.36 | 4.67 | 6.54 | 3.86 | 4.46 |
| FSC threshold | 0.143 | 0.143 | 0.143 | 0.143 | 0.143 | 0.143 | 0.143 | 0.143 |
| **Refinement** | | | | | | | | |
| Initial model used (PDB code) | 3CUE 3PR6 2J3T 2J3W | 3CUE 3PR6 2J3T 2J3W | | 3CUE 3PR6 2J3T 2J3W | 3CUE 3PR6 2J3T 2J3W | 3CUE 3PR6 2J3T 2J3W | 3CUE 3PR6 2J3T 2J3W 3RWO | 3CUE 3PR6 2J3T 2J3W 3RWO |
| Model resolution (Å) | 3.71 | 4.15 | | 6.54 | 4.67 | 4.36 | 3.86 | 4.46 |
| FSC threshold | 0.5 | 0.5 | | 0.5 | 0.5 | 0.5 | 0.5 | 0.5 |
| Map sharpening B factor (Å) | -100 | -180 | | -120 | -120 | -120 | -90 | -155 |
| Model composition | | | | | | | | |
| Non-hydrogen atoms | 23371 | 23082 | | 46742 | 46453 | 46164 | 24944 | 49888 |
| Protein residues | 3292 | 3298 | | 6584 | 6590 | 6596 | 3492 | 6984 |
| Ligands | 0 | 0 | | 0 | 0 | 0 | 0 | 0 |
| B factors (Å$^2$) | | | | | | | | |
| Protein | 92.08 | 121.61 | | 92.44 | 107.24 | 122.3 | 69.39 | 69.48 |
| Ligand | —— | —— | | —— | —— | —— | —— | —— |
| R.m.s.deviations | | | | | | | | |
| Bond lengths (Å) | 0.007 | 0.008 | | 0.007 | 0.008 | 0.008 | 0.008 | 0.008 |
| Bond angles (°) | 1.399 | 1.454 | | 1.400 | 1.427 | 1.454 | 1.45 | 1.447 |
| Validation | | | | | | | | |
| MolProbity Score | 1.88 | 2.14 | | 1.88 | 2.03 | 2.15 | 2.27 | 2.29 |
| Clashcore | 6.72 | 9.41 | | 6.82 | 8.16 | 9.44 | 11.09 | 11.55 |
| Poor rotamers (%) | 0.3 | 0.73 | | 0.3 | 0.51 | 0.73 | 1.42 | 1.42 |
| Ramachandran plot | | | | | | | | |
| Favored (%) | 91.4 | 85.72 | | 91.36 | 88.55 | 85.69 | 88.3 | 88.29 |
| Allowed (%) | 8.6 | 14.25 | | 8.61 | 11.42 | 14.28 | 11.5 | 11.50 |
| Disallowed (%) | 0.00 | 0.03 | | 0.03 | 0.03 | 0.03 | 0.20 | 0.20 |

**Table S2.**

**Summary of model building**

| Molecule | Length | Domain/Region | PDB code | Modeling |
|---|---|---|---|---|
| Trs120 | 1289 | 264-1289 | -- | De novo building |
| Trs130 | 1102 | 250-1084 | -- | De novo building |
| Trs65 | 560 | 212-559 | -- | De novo building |
| Tca17 | 152 | 3-146 | 3PR6 | Rigid docking |
| Trs33 | 268 | 33-263 | 2J3T | Homology modeling |
| Bet3-A | 193 | 8-193 | 3CUE | Rigid docking |
| Bet3-B | 193 | 8-190 | 3CUE | Rigid docking |
| Bet5 | 159 | 2-157 | 3CUE | Rigid docking |
| Trs23 | 219 | 1-219 | 3CUE | Rigid docking |
| Trs31 | 283 | 25-282 | 3CUE | Rigid docking |
| Trs20 | 175 | 2-173 | 2J3W | Homology modeling |
| Ypt32 | 222 | 7-200 | 3RWO | Rigid docking |
